# Ykt6 membrane-to-cytosol cycling regulates exosomal Wnt secretion

**DOI:** 10.1101/485565

**Authors:** Karen Linnemannstöns, Pradhipa Karuna, Leonie Witte, Jeanette Kittel, Adi Danieli, Denise Müller, Lena Nitsch, Mona Honemann-Capito, Ferdinand Grawe, Andreas Wodarz, Julia Christina Gross

## Abstract

Protein trafficking in the secretory pathway, for example the secretion of Wnt proteins, requires tight regulation. These ligands activate Wnt signaling pathways and are crucially involved in development and disease. Wnt is transported to the plasma membrane by its cargo receptor Evi, where Wnt/Evi complexes are endocytosed and sorted onto exosomes for long-range secretion. However, the trafficking steps within the endosomal compartment are not fully understood. The promiscuous SNARE Ykt6 folds into an auto-inhibiting conformation in the cytosol, but a portion associates with membranes by its farnesylated and palmitoylated C-terminus. Here, we demonstrate that membrane detachment of Ykt6 is essential for exosomal Wnt secretion. We identified conserved phosphorylation sites within the SNARE domain of Ykt6, which block Ykt6 cycling from the membrane to the cytosol. In *Drosophila*, Ykt6 RNAi-mediated block of Wg secretion is rescued by wildtype but not phosphomimicking Ykt6. We show that phosphomimicking Ykt6 accumulates at membranes, while wildtype Ykt6 regulates Wnt trafficking between the plasma membrane and multivesicular bodies in a dose-dependent manner. Taken together, we show that exosomal sorting of Wnts is fine-tuned by a regulatory switch in Ykt6 shifting it from a membrane-bound to cytosolic state at the level of endosomal maturation.

## Introduction

Protein trafficking is a dynamic cellular process that fulfills information exchange with the cell’s surrounding by coordinated uptake and release of cargo proteins. Especially the release of signaling proteins is tightly regulated. For example, Wnt proteins are secreted from source cells and act on neighboring target cells to activate Wnt signaling pathways, which play a central role in stem cell maintenance, differentiation in development and adult homeostasis (Nusse & Clevers, 2017). The first trafficking step within the Wnt secretion pathway is the lipidation of Wnts in the ER by Porcupine (Kadowaki *et al*, 1996; Tanaka *et al*, 2000). This modification is required for their activity and secretion and is essential for p24 protein-dependent Wnt exit from the ER (Buechling *et al*, 2011; Port *et al*, 2011). Here, the cargo receptor Evi (also referred to as Wntless) recognizes palmitoleic acid-modified Wnts and escorts them from the ER to the plasma membrane (Herr & Basler, 2012). In the ER, availability of Evi depends on Wnt ligands and is regulated by the ERAD pathway (Glaeser *et al*, 2018). The recycling of Evi from the cell surface to the trans-Golgi network (TGN) enables further transport of newly synthesized Wnts from the TGN to the cell surface. Evi recycling depends on clathrin- and adaptor protein 2 (AP-2)-mediated endocytosis (Gasnereau *et al*, 2011), but also retromer function, because blocking this step leads to a reduction in Wnt secretion (Port *et al*, 2008; Belenkaya *et al*, 2008; Yang *et al*, 2008; Harterink *et al*, 2011; Zhang *et al*, 2011). Wnts have been shown to traffic and spread on extracellular vesicles (EV) such as exosomes (Gross *et al*, 2012; Koles *et al*, 2012; Beckett *et al*,2013; Menck *et al*, 2013), for example in the context of spermatogenesis and nerve regeneration (Koch *et al*, 2015; Tassew *et al*, 2017). In addition, Wnt proteins are also secreted into the extracellular space in other forms, such as cytonemes at the cell surface and it is not understood how trafficking of Wnts is channeled into one of these trafficking routes.

We found that active Wnts are trafficking via endosomal compartments and require the soluble N-ethylmaleimide-sensitive-factor attachment receptor (SNARE) Ykt6 for their secretion on exosomes in *Drosophila* and human cells (Gross *et al*, 2012), however at which level of protein trafficking Ykt6 acts and the underlying molecular mechanism remain to be uncovered.

Ykt6 is an unusual SNARE as it is lacking a transmembrane domain and therefore cycles between cytosol and membranes. Membrane localization depends on the intramolecular interaction of the N-terminal Longin and C-terminal SNARE domains and the presence of a reversible palmitoylation within a CCAIM/CAAX motif. This motif is conserved in several other proteins, such as Ras, MARCKS and rac-1 (Gao *et al*, 2009). Depalmitoylation and release of Ykt6 are needed for its recycling (Dietrich *et al*, 2005) and to circumvent its entry into the endosomal multivesicular body pathway (Meiringer *et al*, 2008).

Ykt6 localizes to different membranes (such as ER, Golgi, endosomal membranes and plasma membrane) and was found in variable SNARE complexes *in vitro*. It has therefore been proposed to function as a membrane stress sensor within the secretory pathway of yeast (Dietrich *et al*, 2004). While a role for Ykt6 in homotypic fusion of ER and vacuolar membranes, retrograde Golgi trafficking in yeast and several steps of autophagosome formation under starvation conditions was demonstrated in human cells (Matsui *et al*, 2018), *Drosophila* fat body (Takáts *et al*, 2018) and yeast (Bas *et al*, 2018; Gao *et al*, 2018), its function in secretion in higher eukaryotes remains unclear. Considering the ability of Ykt6 to adapt to multiple cellular localizations, we here investigate Ykt6 as a candidate to orchestrate selected cargo sorting using the example of Wnt proteins. Combining proximity-dependent proteomics, *in vivo* genetic and *in vitro* biochemical analyses, we found that the membrane to cytosol cycling of Ykt6 has an evolutionary conserved function in Wnt secretion in Drosophila and in human cells.

## Results

### Ykt6 SNARE domain has an evolutionary conserved function in Wnt secretion

The polarized epithelium of *Drosophila* wing imaginal discs (WID) is a model system to study the secretory pathway of Wingless (Wg), the *Drosophila* homologue of Wnt1 (reviewed in (Swarup & Verheyen, 2012; Parchure *et al*, 2017)). RNAi-mediated knockdown of Ykt6 in third instar WIDs strongly reduces extracellular Wg staining (Fig. 1A and (Gross *et al*, 2012)), permanent knockdown is however cell lethal. Potential pleiotropic effects of Ykt6 in the secretory pathway thus prompted us to perform time-controlled RNAi of Ykt6 in the posterior compartment of WIDs to follow the secretion of Wg. This causes intracellular Wg accumulation compared to the anterior control compartment (Fig. 1B), suppresses Wnt target gene expression (Supplementary Fig. 1) and ultimately leads to wing notches in adult flies indicating blocked Wg secretion and consequently Wnt signaling defects (Fig. 1C and (Strigini & Cohen, 2000)). Upon knockdown of Ykt6 in WID, Wg accumulated at the cell periphery, while staining for markers of ER, Golgi, early and late endosomes was unchanged compared to the wildtype side (Fig. 1G). We confirmed the RNAi phenotype using two available loss of function alleles (Haelterman *et al*,2014) (Fig. 1D). These alleles are homozygous lethal, confirming the essential role of Ykt6 as in yeast (McNew *et al*, 1997). As expected from these results, mutant clones in WIDs were small compared to control clones, yet Wg accumulated intracellularly within these clones (Supplementary Fig. 1) as observed for RNAi (Fig. 1B and (Gross *et al*, 2012)).

**Fig.1:**
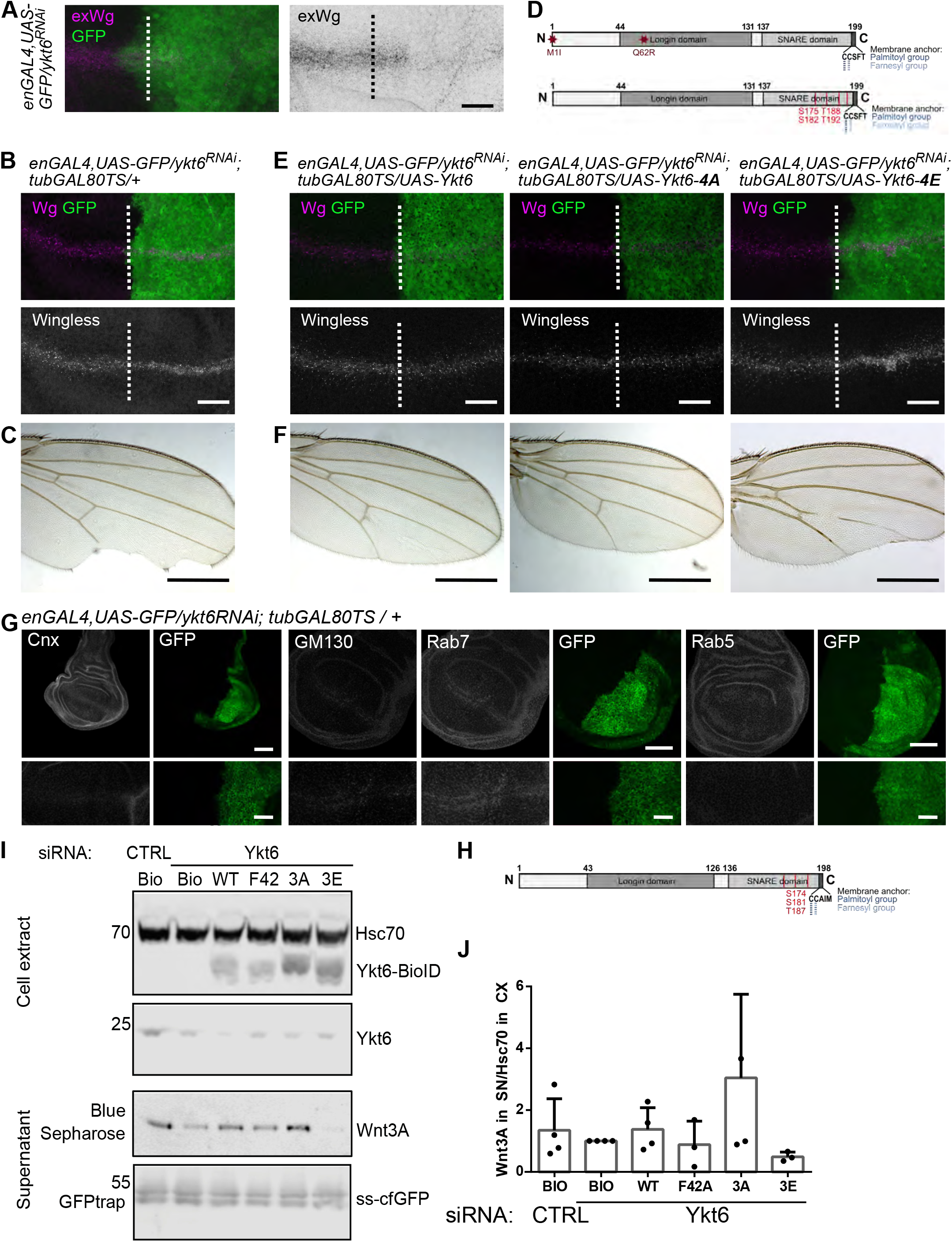
Ykt6 SNARE domain has an evolutionary conserved function in Wnt secretion. (**A**) Knock-down of Ykt6 by RNAi in the posterior compartment of third instar WID marked by co-expression of GFP (engrailed-Gal4, UAS-GFP/UAS-ykt6RNAi) causes extracellular Wingless reduction. (**B, C**) Time-controlled depletion of Ykt6 by RNAi (engrailed-Gal4, UAS-GFP/UAS-ykt6RNAi; tubGal80-TS/+, larvae reared for three days at 29°C) causes intracellular Wg accumulation (**B**) and wing notches (**C**). The GFP-negative anterior compartment serves as an internal control in (A, B). (**D**) Scheme of *Drosophila* Ykt6 mutant alleles (top) and of predicted phosphorylation sites (bottom). (**E, F**) Time-controlled ykt6 RNAi induced block of Wg secretion (**E**) and resulting adult wing margin notches (**F**) can be rescued by cooverexpression of wildtype Ykt6 and non-phosphorylatable Ykt6-4A (left and middle panel), but not by phosphomimetic Ykt6-4E (right panel). Panels in (**A**) show a projection of 12 sections from apical to basal (distance 0,5 μm). Panels in (B) show an overview and a projection of six subapical sections (distance 1 μm). Panels in (E) show an overview and a projection of six subapical sections (distance 0,5 μm). Images are representative of >10 discs from three independent experiments. (**G**) Localization of different organelle markers in wildtype (left side) and Ykt6 RNAi (right side, GFP-positive) from enGAL4, UAS-GFP/Ykt6RNAi WID. Overview (upper panel) and single sections of Calnexin (ER), GM130 (Golgi), Rab5 (EE) and Rab7 (LE). Scale bars represent 500 μm in adult wing images (C,F), 100 μm (B) and 50 μm (E,G) in overview images and 20 μm in all other sections in (B,E,G) and. (**H**) Scheme of human Ykt6 with predicted phosphorylation sites. (**I**) Western Blot analysis of Wnt secretion from Hek293T cells and (**J**) quantification of (**I**), n=3. One-way Anova significance level: n.s.

A recent study described phosphorylation sites within the SNARE domain of nonneuronal SNAREs conserved over the plant, fungi and animal kingdom (Malmersjö *et al*, 2016). These sites are in the SNARE layers facing each other and thereby sterically block the interacting domains of the SNARE helices. As shown for VAMP8, phosphorylation or mutation of these sites inhibits fusion of secretory granules (Malmersjö *et al*, 2016). We analyzed the *Drosophila* Ykt6 sequence and mutated four predicted sites within the SNARE domain to alanine (Ykt6-4A) or glutamic acid (Ykt6-4E) (Fig. 1D). These constructs were expressed by enGal4 in addition to tubGAL80-mediated time-controlled RNAi of Ykt6 in the posterior WID. Block of Wg secretion and wing notches were rescued by expression of wild type and non-phosphorylatable Ykt6-4A (Fig.1E and F, left and middle panel, Supplementary Fig.1), confirming its specificity. However, co-expression of phosphomimicking Ykt6-4E results in Wg accumulation and adult wing defects (Fig.1E and F, right panel, Supplementary Fig.1). Along these lines, Ykt6-WT and to some extent-4A, but not -4E, were able to rescue overall lethality of *ykt6* mutant alleles (Supplementary Fig.1). Within the human Ykt6 sequence we identified and mutated three sites in the SNARE layer to either alanine (Ykt6-3A) or glutamic acid (Ykt6-3E) (Fig. 1H). In Ykt6 RNAi-depleted human Hek293T or Skbr3 cells Wnt secretion is reduced (Supplementary Fig. 2). Endogenous Ykt6 partially localizes in punctae together with Wnt3A, while Ykt6 knockdown causes intracellular accumulation of Wnt3A (Supplementary Fig. 2). Expressing siRNA-resistant, N-terminally-tagged Ykt6 mutant constructs, we found that in contrast to Ykt6-WT and -3A, phosphomimicking Ykt6-3E is unable to rescue Wnt secretion (Fig. 1I and J, Supplementary Fig. 2). Mutation of F42 to alanine, a site within the Longin domain and required for the cytoplasmic, closed conformation of Ykt6 (Tochio *et al*, 2001), did not reduce Wnt secretion. All these constructs did not affect default secretion of secreted GFP (ssGFP (Suzuki *et al*, 2012)) indicating that Ykt6 is specifically involved in the secretion of Wnts (Fig. 1I). This demonstrates an evolutionary conserved function for the Ykt6 SNARE domain in Wnt secretion in *Drosophila* and human, which is impaired by mutation of conserved phosphorylation sites within the SNARE domain.

### Wnt accumulates at post-Golgi membranes together with phosphomimicking Ykt6

We next investigated the intracellular localization of Ykt6 in *Drosophila* and Hek293T cells. Within polarized WID epithelial cells, Ykt6-WT localized to apical, middle and basal planes, while Ykt6-4E mutant staining was increased only basally (Fig. 2A, B). This effect was stronger using N-terminally tagged Ykt6-WT and Ykt6-4E (Fig. 2A, C). In Hek293 cells, overexpressed, tagged Ykt6-WT showed strong staining of the cytoplasm, while Ykt6-3E localization was more punctate at Golgi and plasma membranes (Fig. 2D, Supplementary Fig. 2). To confirm this, we biochemically separated cytosolic and membrane bound proteins by differential detergent fractionation (Baghirova *et al*, 2015). Indeed, Ykt6-3E was enriched in the membrane fraction, while Ykt6-WT was mainly in the cytoplasmic fraction (Fig. 2E and Supplementary Fig. 2). In line with this the majority of endogenous Ykt6 localizes to the cytoplasm (Supplementary Fig. 2 and (Itzhak *et al*, 2016)). This demonstrates that phosphomimicking modifications and therefore putative phosphorylations within the SNARE domain result in a form of Ykt6 that preferentially associates with membranes. We therefore decided to use the phosphomimicking mutant as a tool to dissect the role of Ykt6 in trafficking of Wnts.

**Fig.2:**
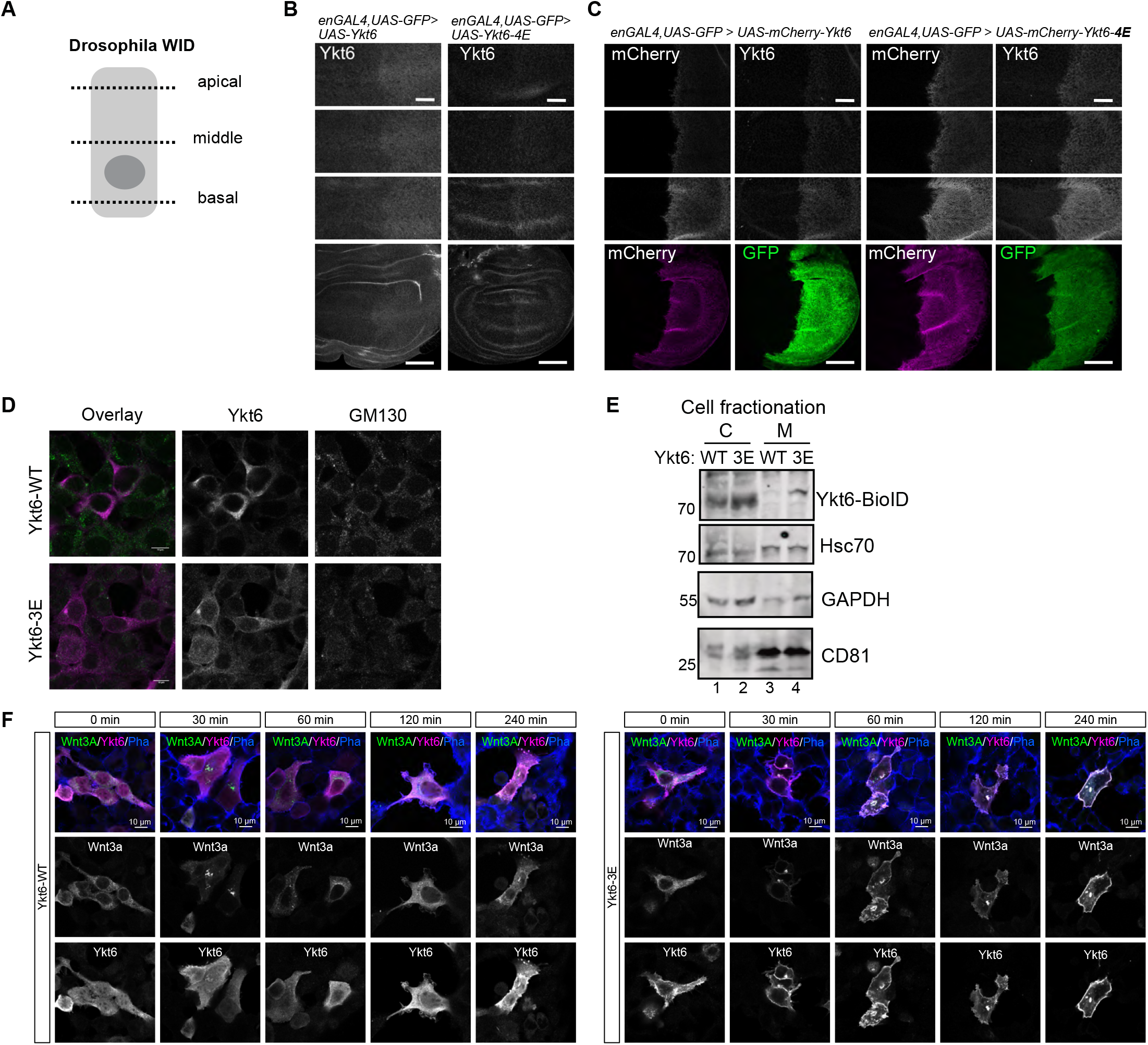
Wnt accumulates at post-Golgi membranes together with phosphomimicking Ykt6. (**A**) Scheme of *Drosophila* WID epithelial cells with apical, middle and basal section represented in (B, C). (**B, C**) Localization of overexpressed untagged (**B**) and mCherry-tagged (**C**) Ykt6-WT (left panels) and Ykt6-4E constructs (right panels). Images are representative of three independent experiments with > six WIDs. Scale bar in (B, C) is 50 μm in overviews and 20 μm in sections. (**D**) Colocalization of N-terminally BioID-tagged Ykt6-WT and -3E in Hek293T cells with Golgi marker GM130. Scale bar in (**D**) is 10 μm and a representative field of view from three independent experiments is depicted. (**E**) Representative blot of detergent fractionation of Hek293 cells transfected with Ykt6-BioID WT and 3E constructs. C - cytoplasmic, M - membrane fraction, n=3. (**F**) Hek293T cells transfected with RUSH-EGFP-Wnt3A in combination with Ykt6-WT (left panel) and Ykt6-3E (right panel) were treated with 50 μM Biotin for the indicated time points and stained for Ykt6 and F-actin in addition to endogenous GFP signal. Scale bar represents 10 μm, representative images of three independent experiments.

To understand the dynamics of Ykt6-dependent Wnt trafficking we used “retention using selective hook” (RUSH), an inducible system for release of secretory cargo (Boncompain *et al*, 2012). This system consists of an ER-resident streptavidin-KDEL fusion protein (“hook”) and a “bait”-protein fused to a streptavidin-binding peptide (SBP) to be retained in the ER. We constructed a tagged Wnt3A with an integrated GFP and an SBP between the signal peptide and the core sequence of Wnt3A. In Hek293T cells this construct is signaling active and localized to ER in the absence of biotin (Supplementary Fig. 3). Addition of 50 μM biotin triggered ER-release of Wnt3A by competitive binding to the streptavidin-hook and 60 to 90 min later Wnt3A localized to the perinuclear region indicating transport to the Golgi (Fig. 2F, Supplementary Fig. 3 and Supplementary Video 1). In cells coexpressing Ykt6-WT, Wnt3A is detected both at the Golgi and in the cytoplasm at later time points (>120 min), partially co-localizing with Ykt6. In contrast to Ykt6-WT, co-expression of Ykt6-3E leads to strong accumulation with Wnt3A at the plasma membrane (>120 min) (Fig. 2F). Hence, phosphomimicking Ykt6 does not block ER to Golgi trafficking of Wnt3A within the secretory pathway but accumulates with Wnt3A at different post-Golgi membranes.

### Ykt6 RNAi defect resembles loss of late secretory pathway SNAREs

Proteins of the SNARE family drive membrane fusion by formation of a trans-SNARE complex consisting of four specific v- and t-SNAREs present at vesicle (v) and target (t) membranes. Albeit some redundancy, different trafficking steps are mediated by specific sets of SNAREs (Dingjan *et al*, 2018). Ykt6 has multiple sites of action and it was shown to interact with different SNARE partners *in vitro* (Tsui & Banfield, 2000). To understand how Ykt6 is involved in post-Golgi Wnt trafficking, we conducted a comparative RNAi candidate approach in *Drosophila* WID, comparing its knockdown to the knockdown of early and late secretory SNAREs. (Fig. 3A, Supplementary Table 1). First, the adult wings of wgGal4-driven RNAi crosses of all 25 SNAREs were analyzed for Wnt signaling defects, i.e. wing notches (Fig. 3A). Due to the general importance of membrane fusion events for protein secretion (Gordon *et al*, 2010), 15 of those 25 SNAREs showed notches and one cross was lethal (Supplementary Table 1). Next, enGal4-driven RNAi of those 16 were analyzed in WID for Wg secretion defects by comparing Wg staining in the anterior with the posterior compartment (Fig. 3A). Under those conditions, six candidates were lethal and ten affected Wg secretion. Golgi SNAREs such as Syx5 and Bet1 strongly reduced Wg secretion and overall cell survival. Ykt6 was recently implicated in autophagosome formation under starvation conditions (Matsui *et al*, 2018; Takáts *et al*, 2018), but under normal growth conditions, autophagic SNAREs such as Syx17 and SNAP29 had no phenotype *in vivo* (Supplementary Table 2). Sec22 and Vamp7 contain a longin domain as Ykt6 and together with Syb act in plasma membrane fusion of secretory vesicles (Gordon *et al*, 2017) and Wg secretion (Li *et al*, 2015; Yamazaki *et al*, 2016; Gao *et al*, 2017). Indeed, we observed Wg accumulation and wing notches for Sec22 and Syb, but not for Vamp7 (Supplementary Table 1). In non-polarized Hek293T reporter cells knockdown of Ykt6 and VAMP1 (human Syb homologue) reduced Wnt activity, while Sec22B and VAMP7 did not (Supplementary Fig. 3). Transverse optical sections clearly showed that Syb RNAi leads to an apical accumulation of Wg like Ykt6 (Fig. 3B). However, staining for Sec22, Syb and Vamp7 in enGAL4/Ykt6-RNAi WID revealed that Ykt6 depletion does not affect localization of these three SNAREs (Supplementary Fig. 3). This indicates that Ykt6 depletion results in a Wg phenotype similar to the late secretory pathway SNARE Syb, a SNARE involved in Wg transcytosis and basolateral secretion in WID (Yamazaki *et al*, 2016).

**Fig.3:**
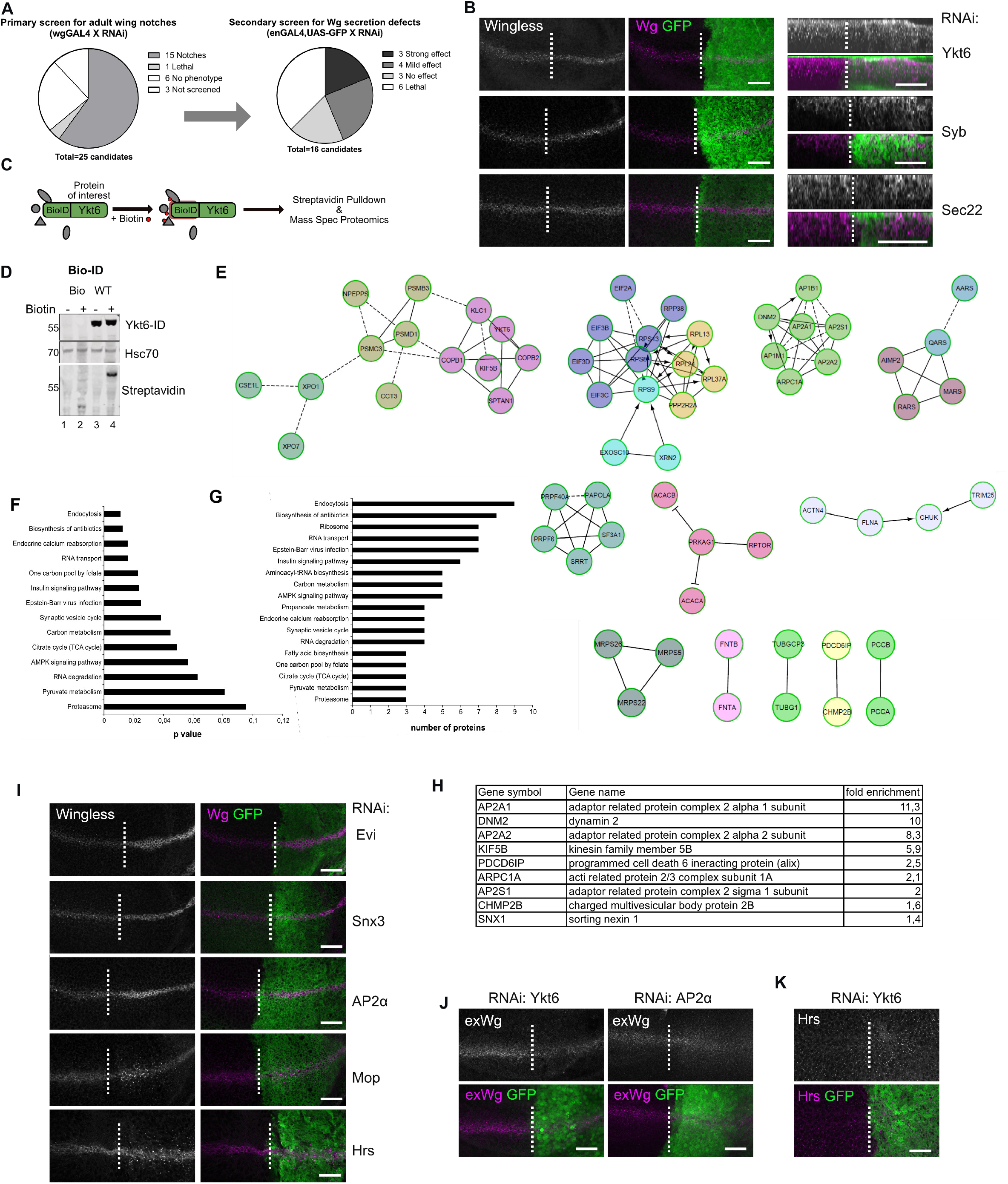
Position of Ykt6 in the secretory pathway. (**A,B**) RNAi against 25 *Drosophila* SNAREs were screened for Wnt secretion defects in adult wings (wgGAL4) and third instar WID (enGAL4), (see also Supplementary Table 1). (**B**) Knockdown of Ykt6 and Syb by RNAi in the posterior compartment of third instar WID marked by coexpression of GFP leads to intracellular Wg accumulation, whereas Sec22 does not affect Wg distribution. The GFP negative compartment serves as an internal control. Panels in (B) show a subapical plane (left) and an optical transverse section (right). (**C**) Schematic of BioID labeling: Ykt6 was N-terminally tagged with a BioID-domain, then upon addition of biotin, streptavidin pull-down was performed and control and Ykt6-WT samples were subjected to proteomics identification. (**D**) Western blot of biotin labeling of Ykt6-BioID and control in the presence of 50 μM biotin. (**E**) Reactome FI network analysis, networks with at least two nodes are displayed, all identified proteins by mass spectrometry in two independent experiments (significance level 0,003 and 1.3-fold over BioID control samples) in Supplementary Table 2. (**F**) Identified proteins were sorted by p-value for Kegg pathway enrichment in Ykt6-WT sample over control identified by BioID approach. (**G**) Number of proteins enriched in different Kegg pathways and (**H**) enrichment score for proteins of the endocytic pathway. (**I**) Wg accumulation phenotypes of different factors required for Wg secretion. RNAi against Evi, Snx3, AP2α, Mop were expressed with enGAL4 and RNAi against Hrs with Dcr; enGAL4. (**J**) RNAi against Ykt6 and AP2α were expressed with enGAL4 and stained for extracellular Wg. (**K**) RNAi against Ykt6 was expressed with enGAL4 and stained for Hrs. Images in (I-K) represent a single confocal section. Images in (B,I-K) are representative of > six WID per RNAi from three independent experiments. Scale bars represent 20 μm.

### Ykt6 acts at the level of endosome maturation

Since Ykt6 does not modulate the localization or expression levels of other SNAREs (Sec22, Vamp7 and Syb) so far described to have a role in Wnt secretion, we envisioned that Ykt6 might fulfill its function independently of a trans-SNARE complex. The subcellular localization of CAAX-motif proteins such as Ras can be rapidly modulated by posttranslational modifications (Salaun *et al*, 2010; Misaki *et al*, 2010). Since Ykt6 also contains a CAAX motif within its sequence, we hypothesized that Ykt6 interaction partners might similarly regulate its cycling between different compartments and could induce location-specific phosphorylation to arrest it at membranes. We used an unbiased BioID approach (Roux *et al*, 2012, 2013) to label proteins in close vicinity of Ykt6 by attaching the prokaryotic BirA* domain N-terminally to Ykt6. This promiscuous ligase biotinylates amine groups of neighboring proteins within a 10 nm radius upon addition of biotin. WT and mock constructs were expressed in the presence of 50μM biotin; biotinylated proteins were purified by streptavidin pulldown and subjected to mass spectrometry (Fig. 3C and D). We identified a total of 143 biotinylated proteins in Ykt6-WT expressing cells enriched over background (Supplementary Table 2). In general, BioID captures weak and transient protein-protein interactions and proximate proteins (Liu *et al*, 2018). Interestingly, we did not identify other SNAREs in the BioID approach. Reactome Functional Network (Wu *et al*, 2010) and Kegg pathway analysis (Kanehisa *et al*, 2016) of identified proteins connected Ykt6 to processes like vesicle trafficking, RNA transport and turnover, metabolic processes and endocytosis (Fig. 3E-G). These connections are in line with the pleiotropic effects observed for Ykt6 in diverse membrane-associated processes like ER-Golgi traffic (McNew *et al*, 1997; Fukasawa *et al*, 2004; Zhang & Hong, 2001), plasma membrane fusion (Gordon *et al*, 2017) and autophagy (Matsui *et al*, 2018; Takáts *et al*, 2018; Bas *et al*, 2018; Gao *et al*, 2018). We found nine candidates that were connected to endocytosis processes (Fig. 3E and H), ranging from early to late endosomal compartments. Knockdown of Dynamin2, Chmp2B and Alix in Hek293T Wnt reporter cells reduced Wnt activity confirming a possible connection between Ykt6 and endosomal sorting in Wnt signaling (Supplementary Fig. 3). We next asked in which step between endocytosis of Wnts from the plasma membrane and sorting into MVBs and onto exosomes Ykt6 is involved. In the polarized epithelium of *Drosophila* WID, Wg is first presented to the apical membrane, then endocytosed for transcytosis and subsequent secretion at the basolateral membrane (Yamazaki *et al*, 2016). The identification of AP2 in the BioID approach and the similarity to the Syb phenotype prompted us to examine depletion of different Wnt secretion components involved after apical plasma membrane presentation of Wg (Fig. 3I). Like Ykt6 and Syb RNAi depletion of AP2α complex components leads to Wg accumulation close to the apical membrane, while knockdown of MVB components HRS and Mop displayed punctate accumulation in Wg secreting and receiving cells (Fig. 3I). Staining for extracellular Wg in non-permeabilized WID was increased in AP2α RNAi, indicating apical surface accumulation of Wg, while extracellular Wg levels were reduced upon loss of Ykt6 (Fig. 3J). Moreover, we observed an increase of Hrs and Lysotracker staining upon Ykt6 RNAi (Fig. 3K, Supplementary Fig. 4), indicating that loss of Ykt6 results in apical intracellular Wg accumulation and affects MVB but not early (Rab5) or late (Rab7) endosomes (Fig. 1G).

### Ykt6 regulates exosome secretion in a concentration-dependent manner

Exosomes are a population of small extracellular vesicles (EVs). They are generated as ILVs by inward budding of the limiting membranes of MVBs and are secreted in an ESCRT-dependent and Alix-Syntenin-regulated manner (Baietti *et al*, 2012). Electron microscopy sections of Ykt6 loss in WID showed no strong morphological defects and MVBs were of similar sizes in WT and Ykt6 RNAi compartments (Supplementary Fig. 4). To understand whether Ykt6 functions at the level of cargo sorting into ILVs we used constitutively active Rab5 to enlarge and visualize endosomes (Stenmark *et al*, 1994). In WID wgGal4-driven Rab5Q88L-YFP expression led to enlarged endosomes positive for endogenous Wg (Fig. 4A). Interestingly, in Ykt6-RNAi WID these were larger and contained less Wg (Fig. 4A, B). In Rab5Q79L-expressing Hek293T cells Wnt3A localization was punctate at the limiting membrane and inside Rab5Q79L-positive endosomes. This pattern was lost upon loss of Ykt6, similarly to loss of the ESCRT machinery by HGS and TSG101 depletion (Supplementary Fig. 4). Overexpressed Ykt6-WT as well as -3E localized inside enlarged endosomes (Fig. 4C), but in Ykt6-3E those endosomes were significantly smaller than in Ykt6-WT (Fig. 4C, D) indicating that the absence of functional Ykt6 enables endosomal fusion events, while the mutated SNARE domain hinders this process. This effect of Ykt6 on endosomal fusion events is clearly aggravated by constitutive Rab5 activity, conceivably because endosomal membrane association of Ykt6 is the rate-limiting step under these conditions.

**Fig.4:**
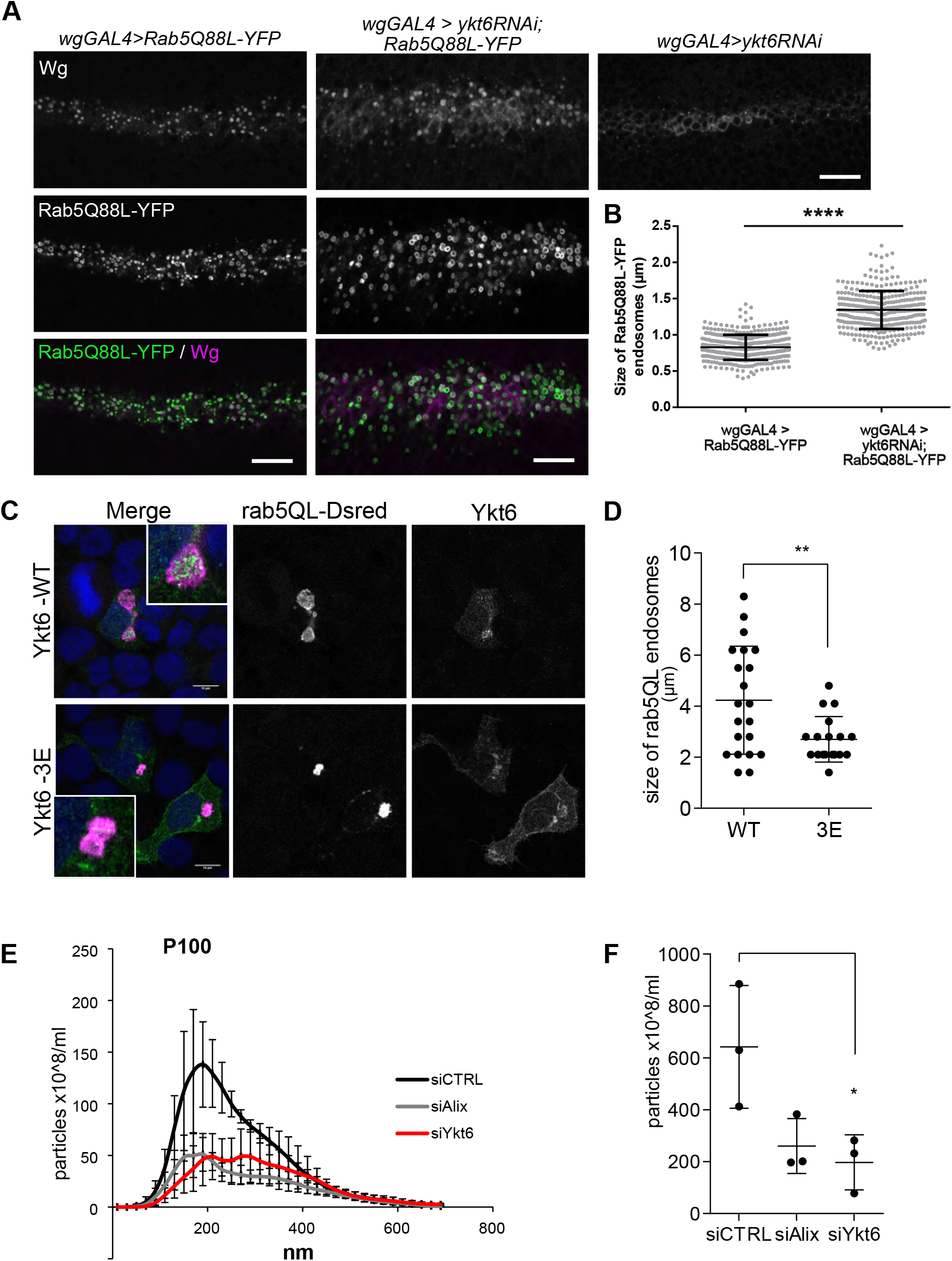
Ykt6 regulates exosome secretion in a concentration-dependent manner. (**A**) Constitutive active Rab5Q88L-YFP was expressed with wgGAL4 alone or in combination with ykt6 RNAi. Images represent a single confocal section; scale bar 10 μm. (**B**) Quantification of (**A**). The diameter of Rab5Q88L-YFP-positive vesicles with a clear lumen was measured. Five representative WIDs from three biological replicates with in total 404 (control) and 326 (ykt6 RNAi) enlarged endosomes were quantified. (**C**) Hek293T cells were co-transfected with plasmids for Rab5Q79L-DsRed and Ykt6-WT or Rab5Q79L-DsRed and Ykt6-3E and analyzed by immunofluorescence microscopy. Scale bars represents 10 μm, n=3. (**D**) Quantification of (C). (**E, F**) Secretion of extracellular vesicles (EV) was analyzed by nanoparticle tracking analysis. Size profile of P100-EV from Hek293T cells transfected with siRNA against control, Ykt6 or Alix. (**F**) quantification of 100-200 nm sized EVs from (E) from three biological replicates.

A portion of ILVs are secreted by fusion of MVBs with the plasma membrane. Thus, next we compared the effect of Ykt6 on EVs secretion into the supernatant of Hek293T cells. In line with the effect of Ykt6 overexpression and RNAi on size of Rab5Q79L endosomes, overexpression of tagged Ykt6-WT significantly reduced secretion of small EVs, but Ykt6-3E did not (Supplementary Fig. 4). Knockdown of Ykt6 as well as Alix reduced the level of EVs as measured by Nanoparticle tracking analysis (NTA) (Fig.4E, F and Supplementary Fig. 4). Thus, recruitment of Ykt6 to endosomal membranes can be a rate-limiting step in exosomes secretion.

These results position Ykt6 function at the level of endosome maturation, where it affects late Wnt trafficking steps for long-range release on exosomes.

### Mutant SNARE domain of Ykt6 blocks closed conformation

In yeast, release of Ykt6 from endosomal membranes into the cytoplasm depends on a functional longin domain and its intramolecular interaction with the SNARE domain to fold into a soluble, closed conformation (Tochio *et al*, 2001) Depalmitoylation of Ykt6 was described to prevent its sorting into MVBs (Meiringer *et al*, 2008). In our BioID approach Ykt6 was highly enriched among the biotinylated proteins (Fig. 3C) indicating an ability of the N-terminal BioID domain to intra- or intermolecularly label Ykt6. Hypothesizing that the closed conformation of Ykt6 would interfere with self-labeling, we compared the biotin-labeling ability of Ykt6-3E with different Ykt6 point mutations known to interfere with the closed conformation, (Fig. 5A). N-terminally BirA*-tagged constructs were expressed in Hek293T cells, treated with biotin, then cytosolic and membrane fractions were analyzed by immuno-blotting for streptavidin-labeled Ykt6 (Fig.5B-D). Compared to Ykt6-WT, the phosphomimicking (3E) and the palmitoyl/farnesyl (C194/195A) double mutant constructs were strongly labeled by the N-terminal BioID domain (Fig.5B, C). Minor self-labeling of Ykt6-WT suggests that a large portion acquires a closed conformation, due to binding of the functional longin domain to the irreversible farnesyl group. In contrast, mutations of both palmitoylation and farnesylation sites (Fukasawa *et al*, 2004) and the phosphomimicking mutations lead to a more unstable and open conformation, allowing stronger self-labeling by the N-terminal BioID (Fig. 5C). Furthermore, protein fractionation confirmed that different Ykt6 mutant constructs vary in their ability to attach to membranes. Whereas Ykt6-WT and -palmitoyl mutant (Ykt6-C194A) were exclusively found cytoplasmic, a substantial portion of Ykt6-3E was attached to membranes (Fig. 5D). To recapitulate these mechanistic insights into Ykt6 membrane association, we analyzed untagged Ykt6 constructs by differential detergent fractionation (Baghirova *et al*, 2015). Here, only Ykt6-3E substantially localized to the membrane fraction (Fig. 5E, F), confirming an additional stabilizing effect of the N-terminal tag as observed in WID (Fig. 2C). In order to further analyze increased Ykt6-3E membrane binding, we monitored the steady-state level of palmitoylated Ykt6-WT and -3E in a click-palmitate assay (Haberkant *et al*, 2016). Ykt6-3E and Wnt3A were both detected in the pull down of all palmitoylated proteins, while Ykt6-WT was below the detection limit (Fig. 5G). This is in line with the notion that Ykt6-WT quickly reverts into its autoinhibited, depalmitoylated form (Fukasawa *et al*, 2004), while depalmitoylation of Ykt6-3E is hindered and therefore a portion remains associated to membranes. This indicates that a functional SNARE domain is required for the conformational changes regulating turnover of palmitoylation and detachment from membranes. Taken together our results strengthen the idea of Ykt6 cytosol to membrane cycling as a switch within the late secretory pathway regulating sorting of selected cargo, such as Wnts, into degradative or secretory pathways.

**Fig.5:**
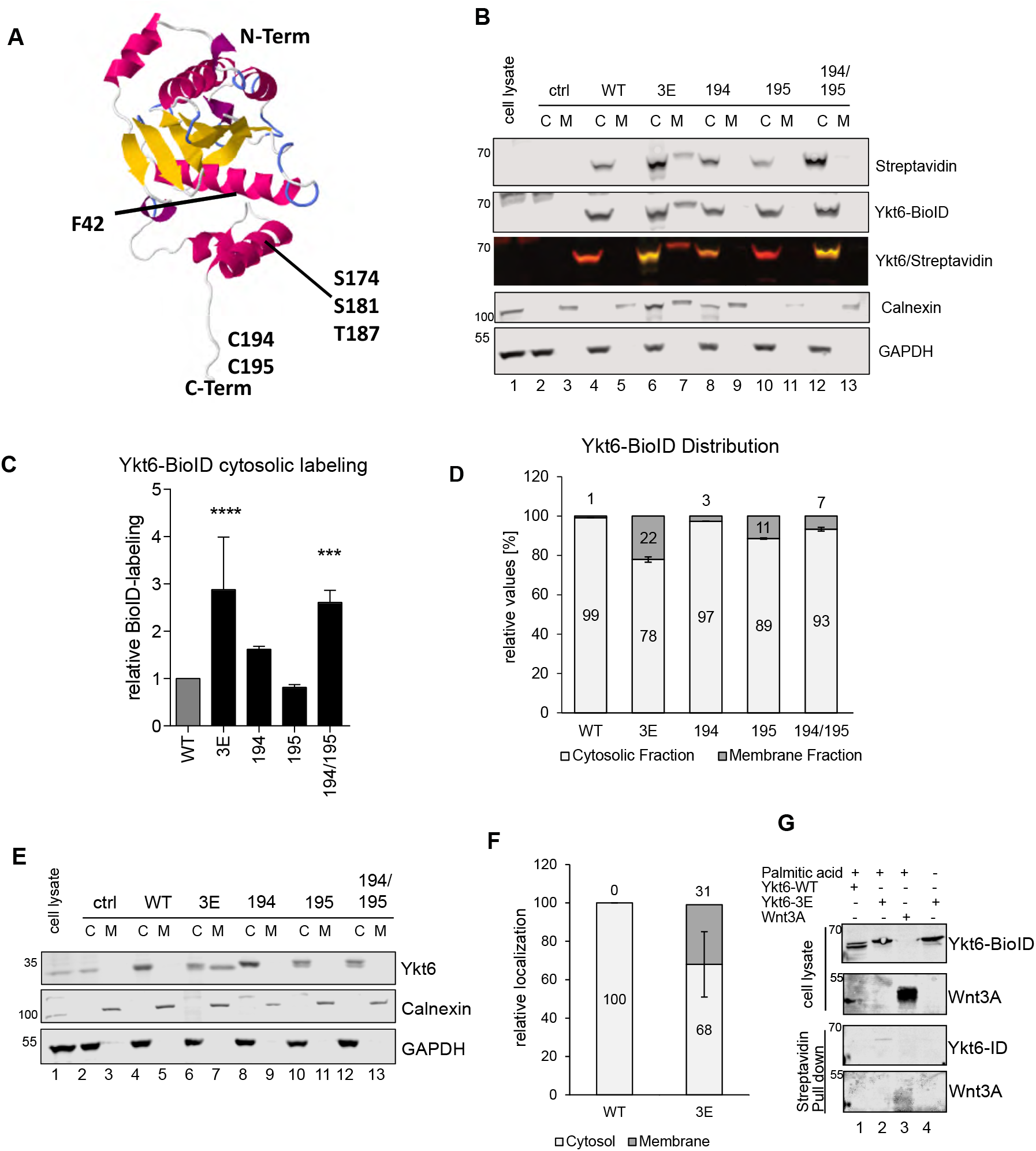
Mutant SNARE domain of Ykt6 blocks closed conformation. (**A**) Structural model of Ykt6 from template (3kyqA). Mutations as indicated: F42 in the Longin domain, S174, S181 and T187 in the SNARE domain and C194 and C195 in the CAAX motif for acylation. (**B**) Biotin-labeling assay. Immunoblot of cell fractionation of different Ykt6-BioID constructs transfected in Hek293T cells, stained with streptavidin, Ykt6 and fraction markers GADPH for cytoplasmic (C) and Calnexin for membrane (M) fraction (n=4). (**C**) Quantification of biotin-labeling in the cytosolic fraction of (**B**). (**D**) Quantification of Ykt6 distribution into cytosolic and membrane fraction from (**B**). (**E**) Immunoblot of cell fractionation of different untagged Ykt6 mutant constructs transfected in Hek293T cells and stained with Ykt6 and fraction markers as in (B) (n=3). (**F**) Quantification of membrane association of Ykt6-WT and - 3E from (E). (**G**) Click palmitoylation assay of Ykt6-WT and -3E, Wnt3A as a positive control. Representative immunoblot of three biological replicates.

## Discussion

Peripheral membrane proteins have an advantage over transmembrane proteins in that their subcellular localization can be rapidly modulated. Here, we demonstrate that the function of Ykt6 for Wnt secretion is tightly linked to its ability to cycle between cytosol and membranes. As shown by NMR structure studies Ykt6 undergoes conformational changes at the interface between its longin domain and the SNARE core (Wen *et al*, 2010). Within the SNARE domain of Ykt6, we identified putative phosphorylation sites, conserved in non-neuronal SNAREs that allow membrane stabilization and prevent depalmitoylation. These identified phosphorylation sites are directly facing the longin domain in crystal structure of the closed conformation (Tochio *et al*, 2001). Our observations for Ykt6-4E/-3E mutants are in line with impaired cycling of mutant Ykt6F42S resulting in permanent membrane localization (Meiringer *et al*, 2008). The phosphorylation sites seem to be widely conserved in non-neuronal SNAREs, as phosphorylation of VAMP8 leads to a block of IgE secretion, by structurally preventing membrane SNARE fusion after docking (Malmersjö *et al*, 2016). The role of Ykt6 in membrane fusion is less clear, as previous studies suggest that Ykt6 can substitute other/multiple SNARE in fusion reactions *in vitro* and under loss of function conditions (Tsui & Banfield, 2000), but in yeast Ykt6 is not directly required for vacuole membrane fusion but rather for docking (Meiringer *et al*, 2008). This agrees with our results and supports a model in which Ykt6 could serve as an adaptor in different membrane/organelles settings. Our unbiased BioID approach positions Ykt6 function at the cross section of metabolic processes and the secretory pathway-at the level of endosomes. Based on our results, we propose a model for Ykt6 to function as a membrane sorting sensor that gets associated with membranes when cargo/cargo receptors accumulate and decides in which trafficking pathway to channel those cargo. BioID approach also identified other processes related to RNA stability and metabolism. Further work will be required to provide functional insight into potential signaling networks activating Ykt6 membrane association.

Recruitment of Ykt6 to endosomal membranes might depend on posttranslational modifications within its SNARE domain and differential binding to interaction partners to decide between secretion and autophagic degradation pathways. Our data indicates that Ykt6 is proximal of very different cellular processes such as endocytosis, RNA transport and metabolic signaling pathways. Interestingly, we did not find any other SNAREs in proximity of Ykt6 under normal conditions. Three recent studies implicated Ykt6 in autophagosome formation under starvation conditions (Kimura *et al*, 2017; Matsui *et al*, 2018; Takáts *et al*, 2018) and another longin SNARE Sec22b mediates unconventional secretion of cytosolic proteins via secretory autophagy (Kimura *et al*, 2017). Interestingly, Ykt6 depletion reduced the number of secreted EVs only under normal condition, while the numbers of secreted EVs from serum-starved cells were similar (Supplementary Fig. 4). Here, we mainly investigated Ykt6 function under normal growth conditions as Wnt secretion is strongly reduced under starvation (Mihara *et al*, 2016). In line with its proposed role as a stress sensor in yeast (Dietrich *et al*, 2004), the majority of human Ykt6 localizes cytoplasmic, potentially serving as a reserve pool to release trafficking stress at different levels and under specific circumstances. In the polarized epithelium of *Drosophila* WID phosphomimicking Ykt6 blocks Wnt secretion similarly to Ykt6 depletion. What could stimulate Ykt6 phosphorylation and therefore association with membranes? Prediction according to NetPhos 3.1 (Blom *et al*, 2004) suggested that the identified phosphorylation sites are potential PKC sites (Supplementary Fig. 4). As PKC gets activated in the context of endocytosis to recruit adaptor complexes to endosomes (Lau *et al*, 2010; Nazarewicz *et al*, 2011), local PKC-dependent conformation change and subsequent palmitoylation could stimulate Ykt6 association with endosomes, yet the physiological relevance of Ykt6 phosphorylation remains to be demonstrated. In general, members of the SNARE family are regulated by posttranslational modifications such as monoubiquitination (Syx5) (Huang *et al*, 2016) and palmitoylation (SNAP25) (Gonzalo & Linder, 1998).

A recent study revealed that Wnt secretion is controlled by the ER stress machinery. Boutros and colleagues demonstrated that in the absence of Wnts its trafficking receptor Evi is destabilized by VCP complex and ubiquitination. Thus Wnt secretion is regulated already early in the secretory pathway by availability of its trafficking receptor in a “demand-and-supply” fashion (Glaeser *et al*, 2018). Our data confirm and extend the notion that Wnt secretion is fine-tuned at several levels of the secretory pathway. We propose that Ykt6 serves as a switch for endosomal pressure release - fine-tuning Wnt secretion in the late secretory pathway and integrating different upstream signaling pathways via a conformational switch in its SNARE domain. Within the polarized epithelium of *Drosophila* WID, apically presented and subsequently endocytosed Wg might serve as a signaling reservoir that can get rapidly mobilized by Ykt6-mediated secretion at the basolateral side. In lines with this idea is a recent study demonstrating that Wg is endocytosed apically, while its receptor Fz2 is internalized from the basolateral side of WID and both meet in acidified endosomes for signal transduction (Hemalatha *et al*, 2016). Moreover, Wg endocytosis from the apical side seems to depend on HSPGs, which are also involved in cargo sorting onto exosomes via Alix and syntenin (Ghossoub *et al*, 2014). Further investigation is required to understand the regulatory networks upstream of Ykt6 membrane recruitment at the crossroad of exosomal secretion and autophagosome induction. Yet with its ability to adapt to multiple cellular localizations, Ykt6 is an ideal candidate to orchestrate selected cargo sorting in the endosomal system.

## Material and Methods

### Plasmids and siRNA

The coding region of *Drosophila* Ykt6 was amplified and the PCR product recombined into pDONR^TM^221 vector using the Gateway BP Clonase II Enzyme mix (Life Technologies, Carlsbad, CA, US). Point mutations of potential phosphorylation sites (S175, S182, T188, T192) were introduced by site directed mutagenesis. For generation of transgenic flies, constructs were suncloned into expression vectors pUASt-attB-rfA-mCherry and pUASt-attB-mCherry-rfA (kind gift from Sven Bogdan) by LR recombination (Life Technologies, Carlsbad, CA, US). Human Ykt6 was amplified from hYkt6-Myc (C-Terminal myc-destination plasmids (DKFZ – Genomics and Proteomics Core Facility)) and the PCR product inserted into pcDNA3.1MycBioID (Addgene #35700). Point mutations for Ykt6-3A (S174A, T181A, S187A), Ykt6-3E (S174E, T187E, S181E), F42A, C194A, C195A and relevant combinations were introduced by site directed mutagenesis. MycBioID tag was removed via Nhe1/Xho1 to obtain untagged constructs in pcDNA3.1. RUSH-EGFP-Wnt3A was constructed by amplifying the core protein sequence of Wnt3A and integrating it by Gibson cloning(Gibson *et al*, 2009) with the Wnt3A signal peptide and the streptavidin-binding peptide sequence into the ER-hook containing pCMV-KDEL-IVS-IRES-reporter plasmid backbone(Boncompain *et al*, 2012). The following expression constructs were used: TCF4/Wnt-Firefly Luciferase(Demir *et al*, 2013), Actin-Renilla Luciferase(Nickles *et al*, 2012), pCMV-Wnt3A(Gross *et al*, 2012), DsRed-Rab5-QL (E. De Robertis, Addgene #29688) and ss-cf-sGFP2 (I. Wada, Addgene #37535).

Dharmacon siRNA SMARTpools were used against:

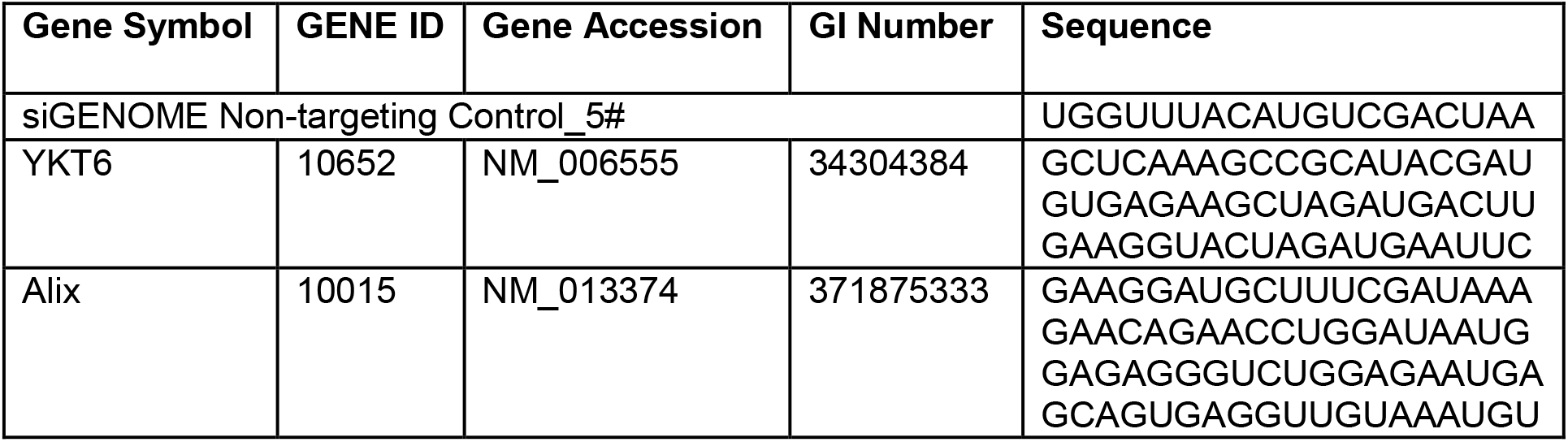

### Antibodies

Antibodies were used against Calnexin, 1:1,000 (WB; rabbit (sc-11397), Santa Cruz), and 1:10 (IF; mouse (Cnx99A 6-2-1), DSHB), CD81 (1.3.3.22), 1:1,000 (WB; mouse (DLN-09707), Dianova), EEA1, 1:300 (IF; mouse (610456), BD), GAPDH (6C5), 1:5,000 (WB; mouse (AM4300), Ambion), GFP 1:1000 (IF; mouse (A11120) and rabbit (A11122), Molecular Probes), GM130, 1:300 (IF; mouse (610823), BD), GM130 1:500 (IF; rabbit (ab30637) Abcam), Hrs 1:10 (IF; mouse (Hrs8-2 and Hrs27-4), DSHB), Hsc70 1:2000 (WB; mouse (sc-7298), Santa Cruz), Lamp1 1:100 (IF; rabbit (ab30687), Abcam), Lamp1 1:250 (IF; rabbit (ab24170), Abcam), mCherry 1:1000 (IF; rabbit (ab167453), Abcam), Rab7 1:10 (mouse (Rab7), DSHB), Sec22, Syb and Vamp7 (IF; 1:250, were kind gifts from Andrew A. Peden(Gordon *et al*, 2017)), Sens (IF; rabbit 1:1000, a kind gift from Hugo Bellen(Nolo *et al*, 2000)), Syx1A 1:10 (mouse (8C3), DSHB), TSG101, 1:1,000 (WB; rabbit (HPA006161), Sigma), Wg, 1:3 for extracellular and 1:20 for total staining (mouse, 4D4, DSHB), Wnt3A, 1:500 (WB; rabbit, Abcam), Wnt5A, 1:500 (WB; rabbit (2530), CST), Ykt6 (WB and IF; mouse (sc-365732), Santa Cruz). Antibodies against Ykt6 were generated by immunizing two guinea pigs with the peptides KVSADQWPNGTEATI (aa 105-119, within Longin domain) and YQNPVEADPLTKMQN (aa 131-145, covers part of SNARE domain). Final bleeds were pooled and affinity purified against the original peptides (Eurogentec). Secondary antibodies directed against the species of interest were coupled to Alexa Fluor 488, 568, 594 and 647 (IF, 1:500, Invitrogen) and 680RD and 800CW (WB, 1:20,000, LiCor).

### Drosophila stocks and genetics

The following Drosophila stocks were used in this study: *en-GAL4,UAS-GFP* (chr. II, a gift from J. Grosshans), *wg-Gal4* (chr. II, a gift from S. Cohen). The following stocks were obtained from Bloomington Drosophila stock center: *da-GAL4* (#5460), *UAS-Dcr; enGAL4,UAS-GFP* (#25752), *tub-GAL80TS* (#7108), *ykt6^A^FRT19A / FM7c,Kr-GAL4,UAS-GFP* (#57143), *ykt6^C^FRT19A / FM7c,Kr-GAL4,UAS-GFP* (#57142), *His2Av-GFP,hsFlp,FRT19A* (#32045), FRT19A (#1709) and *vas-PhiC31; attP.ZH-86Fb* (#24749), AliX TRiP (#33417), Hrs TRiP (# 28026 and #33900). The following UAS-RNAi stocks were obtained from Vienna Drosophila RNAi Center: ALiX (GD32047), AP-2α (GD15565), Evi (GD5214, KK 103812), Mop (KK104860), Sec22 (KK100766), Snx3 (KK104494), Syb (KK102922), Vamp7 (KK108733) and Ykt6 (KK105648). UAS-Ykt6 transgenic lines were generated according to standard protocols by φC31 integrase-mediated site-specific insertion in the attP landing site at ZH-86Fb(Bischof *et al*, 2007). Fly stocks were kept on standard medium containing agar, yeast and corn flour. Crosses were performed at 25°C except for *tub-Gal80TS* crosses, which were moved to 29°C three days before dissection of wing imaginal discs. To generate negatively marked *ykt6* mutant and *FRT* control clones in the wing imaginal disc under the control of *hsFlp*, animals of the appropriate genotype were heat-shocked four days after egg laying for 2h at 37°C on two consecutive days and dissected on the next day in the wandering L3 stage.

### Cell culture and Transfection

Hek293T, HCT116 and SkBr3 cells were maintained in DMEM (Gibco) supplemented with 10% fetal calf serum (Biochrom) at 37 °C in a humidified atmosphere with 5% CO2. Cells were transiently transfected with Screenfect siRNA for siRNA and Screenfect A (Screenfect) for plasmids according to the manufacturer’s instructions and checked regularly for mycoplasma contamination.

### Blue Sepharose Precipitation

The relative amount of Wnts secreted into cell culture supernatant was analyzed using Blue Sepharose precipitation as described(Willert *et al*, 2003; Glaeser *et al*, 2016). Shortly, HEK293T cells were transiently transfected in 6-well plates with 1 μg of Wnt3A plasmids. 72 h after transfection the supernatant was collected and centrifuged at 4000 rpm to remove cell debris, transferred to a fresh tube and rotated at 4°C for 1 h with 1 % Triton X-100 and 40 μl of Blue Sepharose beads. The samples were washed and eluted from the beads using 2X SDS buffer with β-mercaptoethanol and analyzed by immunoblotting.

### Extracellular Vesicles purification

Extracellular Vesicles were purified by differential centrifugation as described previously(Théry *et al*, 2006; Menck *et al*, 2017). In short, supernatants from mammalian cells were subjected to sequential centrifugation steps of 750g, 1500g and 14,000g, before pelleting exosomes at 100,000g in a SW41Ti swinging bucket rotor for 2h (Beckman). Supernatant was discarded and exosomes were taken up in 1/100 of their original volume in H2O.

### Immunostainings, microscopy and image analysis

For immunofluorescence, cells were reverse transfected with siRNAs, seeded in 6 well dishes or 8 well microscopic coverslips, 24h later transfected with indicated plasmids, and 48-72h later fixed with 4% paraformaldehyde. Cells were permeabilized with 0.1% Triton X-100 and blocked in 10% BSA/PBS. Primary antibodies in PBS were incubated for 1h at room temperature and antibody binding visualized by fluorochrome-conjugated secondary antibodies.

Immunostainings of wing imaginal discs were performed as per standard procedures. Total and extracellular Wg staining was carried out as previously described^27^. Staining and microscopy conditions were kept identical for discs used for comparisons. Imaginal discs were mounted in Mowiol and images were taken using a Zeiss LSM780 confocal microscope. Z stacks were generated with 0.5-1μm intervals using a Plan Neofluar 63X/oil NA 1.4 objective. Confocal images were processed with Zen lite (Zeiss), Fiji/ImageJ (NIH)(Schneider *et al*, 2012; Schindelin *et al*, 2012; Rueden *et al*, 2017) and Affinity Designer (Affinity).

The time-lapse movie of RUSH-EGFP-WNT3A was recorded with a Zeiss LSM780 confocal microscope equipped with a LCI Plan Neofluar 63X/water NA 1.3 objective at 37°C. The Z stack was obtained with a frame size of 256x256 px (pixel size 0,24 μm x 0,24 μm) at an interval of 60 sec and consisted of 58 slices with 0,63 μm intervals. Maximum intensity projection over time was generated with Zen lite (Zeiss).

### Electron microscopy

Wing imaginal discs were fixed in 2.5% glutaraldehyde in 100 mM phosphate buffer pH 7.2, washed in 100 mM phosphate buffer and postfixed in 2% osminum tetroxide in phosphate buffer for 1 hour on ice. After contrasting en bloc in 2% uranyl acetate, the specimens were dehydrated in EtOH and embedded in araldite using acetone as an intermediate solvent. Thin sections were stained with 2% uranyl acetate and lead citrate. Sections were observed under an EM 109 (Zeiss) microscope at 80 KV. Quantification of MVB diameter was done manually in Fiji/ImageJ (NIH)(Schneider *et al*, 2012; Rueden *et al*, 2017; Schindelin *et al*, 2012).

### Click Palmitoylation Assay

Click assay was performed as described previously(Haberkant *et al*, 2016). In short, HEK293T cells were seeded, then transfected with plasmids (YKT6-WT BioID, YKT6-3E BioID, Wnt3A) in DMEM supplemented with 10% FBS. ω-alkynyl palmitic acid (Alk-C16) was dissolved in Ethanol to a final concentration of 50mM and stored at −80°C. Alk-C16 was diluted to a final concentration of 100 μM in DMEM supplemented with 5% FBS (fatty acid-free) sonicated for 15 min at room temperature in a water bath and then allowed to precomplex for another 15 min. Alk-C16-containing medium was added to cells and partially replaced after 24 hours. 72 hours post transfection, cells were lysed (PBS with 1% Triton x-100, 0.1% SDS, PIC), then centrifuged at 16,000g for 5 min at 4°C. Then lysates were precipitated with Wessel-Flugge Protein precipitation. The click labelling reaction (0.1 mM biotin-azide, 1 mM Tris(2-carboxyethyl)phosphine hydrochloride (TCEP, Sigma-Aldrich) dissolved in water, 0.1 mM Tris[(1-benzyl-1H-1,2,3-triazol-4-yl)methyl]amine (TBTA, Sigma-Aldrich) dissolved in DMSO and 1 mM CuSO4 in water) was incubated shaking for 2 hours at 37°C under dark conditions. After the click reaction, the samples were precipitated with 10x MeOH overnight at −80° C, then centrifuged and washed again with ice cold Methanol, The dried pellet was resuspended in 4% SDS. Click-biotinylated proteins precipitated with High Capacity Neutravidin Agarose Resin (Thermo Scientific). Samples were washed with 1% SDS and eluted, then analyzed further by immunoblotting.

### BioID pull down & Mass spectrometry

For large-scale BioID pull down, cells were seeded and 24hours later transfected with BioID-WT or mock constructs. 36 hours post transfection 50 μM biotin was added over night. Cells were washed with PBS twice and harvest in Ripa Lysis buffer (50mM Tris-HCl pH7.5, 150mM NaCl, 1% Igepal, 0.5% Sodiumdesoxycholate, 0.1% SDS) containing 1× Complete protease inhibitor (Life Technologies). After centrifugation at 16,500 × g for 10 min, lysates were boiled 5 min in non-reducing SDS sample buffer (300mM Tris-HCl pH 6.8, 12% SDS, 0.05% Bromphenolblue, 60% Glycerol, 12mM EDTA), either fully separated or run short-distance (1.5 cm) on a 4-12 % NuPAGE Novex Bis-Tris Minigel (Invitrogen). Gels were stained with Coomassie Blue for visualization purposes. Full lanes were sliced into 23 equidistant slices regardless of staining, short runs cut out as a whole and diced. After washing, gel slices were reduced with dithiothreitol (DTT), alkylated with 2-iodoacetamide and digested with trypsin overnight. The resulting peptide mixtures were then extracted, dried in a SpeedVac, reconstituted in 2% acetonitrile/0.1% formic acid/ (v:v) and prepared for nanoLC-MS/MS as described previously(Atanassov & Urlaub, 2013).

For generation of a peptide library for SWATH-MS, equal amount aliquots from each sample were pooled to a total amount of 80 μg, and separated into eight fractions using a reversed phase spin column (Pierce High pH Reversed-Phase Peptide Fractionation Kit, Thermo Fisher Scientific). MS analysis Protein digests were separated by nanoflow chromatography. 25% of gel slices or 1 μg aliquots of digested protein were enriched on a self-packed precolumn (0.15 mm ID x 20 mm, Reprosil-Pur120 C18-AQ 5 μm, Dr. Maisch, Ammerbuch-Entringen, Germany) and separated on an analytical RP-C18 column (0.075 mm ID x 250 mm, Reprosil-Pur 120 C18-AQ, 3 μm, Dr. Maisch) using a 30 to 90 min linear gradient of 5-35 % acetonitrile/0.1% formic acid (v:v) at 300 nl min-1.

For Spectral Counting analysis, the eluent was analyzed on a Q Exactive hybrid quadrupole/orbitrap mass spectrometer (ThermoFisher Scientific, Dreieich, Germany) equipped with a FlexIon nanoSpray source and operated under Excalibur 2.4 software using a data-dependent acquisition method. Each experimental cycle was of the following form: one full MS scan across the 350-1600 m/z range was acquired at a resolution setting of 70,000 FWHM, and AGC target of 1*10e6 and a maximum fill time of 60 ms. Up to the 12 most abundant peptide precursors of charge states 2 to 5 above a 2*10e4 intensity threshold were then sequentially isolated at 2.0 FWHM isolation width, fragmented with nitrogen at a normalized collision energy setting of 25%, and the resulting product ion spectra recorded at a resolution setting of 17,500 FWHM, and AGC target of 2*10e5 and a maximum fill time of 60 ms. Selected precursor m/z values were then excluded for the following 15 s. Two technical replicates per sample were acquired.

SWATH-MS library generation was performed on a hybrid triple quadrupole-TOF mass spectrometer (TripleTOF 5600+) equipped with a Nanospray III ion source (Ionspray Voltage 2400 V, Interface Heater Temperature 150°C, Sheath Gas Setting 12) and controlled by Analyst TF 1.7.1 software build 1163 (all AB Sciex), using a Top30 data-dependent acquisition method with an MS survey scan of m/z 380-1250 accumulated for 250 ms at a resolution of 35 000 full width at half maximum (FWHM). MS/MS scans of m/z 180-1500 were accumulated for 100 ms at a resolution of 17,500 FWHM and a precursor isolation width of 0.7 FWHM, resulting in a total cycle time of 3.4 s. Precursors above a threshold MS intensity of 200 cps with charge states 2+, 3+, and 4+ were selected for MS/MS, the dynamic exclusion time was set to 15 s. MS/MS activation was achieved by CID using nitrogen as a collision gas and the manufacturer’s default rolling collision energy settings. Two technical replicates per reversed phase fraction were analysed to construct a spectral library.

For quantitative SWATH analysis, MS/MS data were acquired using 100 variable size windows(Zhang *et al*, 2015) across the 400-1200 m/z range. Fragments were produced using rolling collision energy settings for charge state 2+, and fragments acquired over an m/z range of 180-1500 for 40 ms per segment. Including a 250 ms survey scan this resulted in an overall cycle time of 4.3 s. Two replicate injections were acquired for each biological sample.

### Mass Spectrometry Data processing

For Spectral Counting analysis, peaklists were extracted from the raw data using Raw2MSMS software v1.17 (Max Planck Institute for Biochemistry, Martinsried, Germany) Protein identification was achieved using MASCOT 2.5.1 software (Matrixscience, London, United Kingdom). Proteins were identified against the UniProtKB Homo sapiens reference proteome (revision 02-2017, 92,928 entries). The search was performed with trypsin as enzyme and iodoacetamide as cysteine blocking agent. Up to two missed tryptic cleavages and methionine oxidation as a variable modification were allowed for. Search tolerances were set to 10 ppm for the precursor mass and 0.05 Da for-fragment masses. Scaffold software version 4.4.1.1 (Proteome Software Inc., Portland, OR) was used to validate MS/MS based peptide and protein identifications. Protein and peptide identifications were filtered to 1% FDR using a concatenated forward-and-reverse decoy database approach. Relative quantification of proteins in the samples was achieved by two-sides t-tests of normalized Spectral Counts using a Benjamini-Hochberg-corrected p value of 0.05 to judge significance. To allow for the calculation of low abundance protein ratios, a minimum value of 3 spectral counts was introduced where necessary to avoid division by zero issues.

For SWATH-MS analysis, protein identification was achieved using ProteinPilot Software version 5.0 build 4769 (AB Sciex) at “thorough” settings. MS/MS spectra from the combined qualitative analyses were searched against the UniProtKB Homo sapiens reference proteome (revision 02-2017, 92,928 entries) augmented with a set of 51 known common laboratory contaminants to identify 597 proteins at a False Discovery Rate (FDR) of 1%. Spectral library generation and SWATH peak extraction were achieved in PeakView Software version 2.1 build 11041 (AB Sciex) using the SWATH quantitation microApp version 2.0 build 2003. Following retention time correction on endogenous peptides spanning the entire retention time range, peak areas were extracted using information from the MS/MS library at an FDR of 1%(Lambert *et al*, 2013). The resulting peak areas were summed to peptide and protein area values, which were used for further statistical analysis. Reactome Functional Network analysis (Wu *et al*, 2010) was performed with Cytoscape (www.cytoscape.org) and Kegg pathway analysis was performed with David(Huang *et al*, 2009, 2008).

### Immunoblot

To analyze total cell lysates using immunoblot, cells were lysed in SDS-PAGE sample buffer, boiled for 5 min. Proteins were separated on 4-12% gradient gels (Bolt Bis-Tris Plus Gels, Thermo Scientific) and transferred to PVDF membrane (Merck). After blocking with 5% (wt/vol) milk-TBST, membranes were incubated with Licor-800nm-conjugated streptavidin (1:20,000, ab7403; Abcam) for 30 min. After detecting biotinylated proteins, membranes were subjected to detection with antibodies against Ykt6 and cellular fraction markers as mentioned and Licor680nm-conjugated secondary antibodies.

### Nanoparticles Tracking Analysis

EV samples were dilute 1:25 in PBS (16μl sample and 384μl PBS). The 400μl were injected to the measuring cell of the NanoSight LM10 machine (Malvern Instruments) and 60s videos were recorded with the NanoSight NTA 2.3 Analytical Software and the sCMOS camera (Shutter 30.01ms, Gain 500m, Frame rate 24.99fps and temperature 22.5°C). A total of five videos were recorded per sample when the measured location was slightly changed in between, to represent the whole sample. The videos were analyzed by the NanoSight NTA 2.3 Analytical Software and the particles concentration, size distribution and the general mean and mode of the samples were obtained.

### Ykt6 model prediction

Ykt6-3E structural model was predicted using RaptorX(Källberg *et al*, 2012) and is based on the Ykt6 structure (3kyqA) as a template.

### Statistics

All experiments were carried out at least in biological triplicates. Error bars indicate s.d. Statistical significance was calculated by carrying out One-way Anova or Student’s t-test where appropriate.

The data that support the findings of this study are available from the corresponding author upon reasonable request.

## Acknowledgements

The authors thank the Core Facility Proteomics at the Institute of Clinical Chemistry, UMG, Jörg Großhans, Hugo Bellen, Sven Bogdan and Andrew Peden for fly reagents.; Thomas Monecke for helpful comments regarding Ykt6 structural model. We thank Varun Chaudhary and Dolma Choezom for critical reading of the manuscript. Research in the lab of JCG is supported by the DFG-funded Research Center SFB1324/1 and GR4810/2-1, the Research program of the University Medical Center, University of Göttingen and a postdoctoral fellowship to K.L. by the Dorothea Schlözer Program, University of Göttingen.

## Author Contributions

K.L., designed and carried out Drosophila experiments and data analysis with the help of J.K and L.N., L.W. P.K., A.D., D.M., M.H.C. and J.C.G. carried out cell culture experiments and data analysis. M.H.C. generated fly lines and constructs. F.G. and A.W. performed electron microscopy analysis. J.C.G. conceived and supervised the study and wrote the paper with the help and comments of all authors.

## Corresponding author

Correspondence to Julia Christina Gross

---

**Fig. S1:**
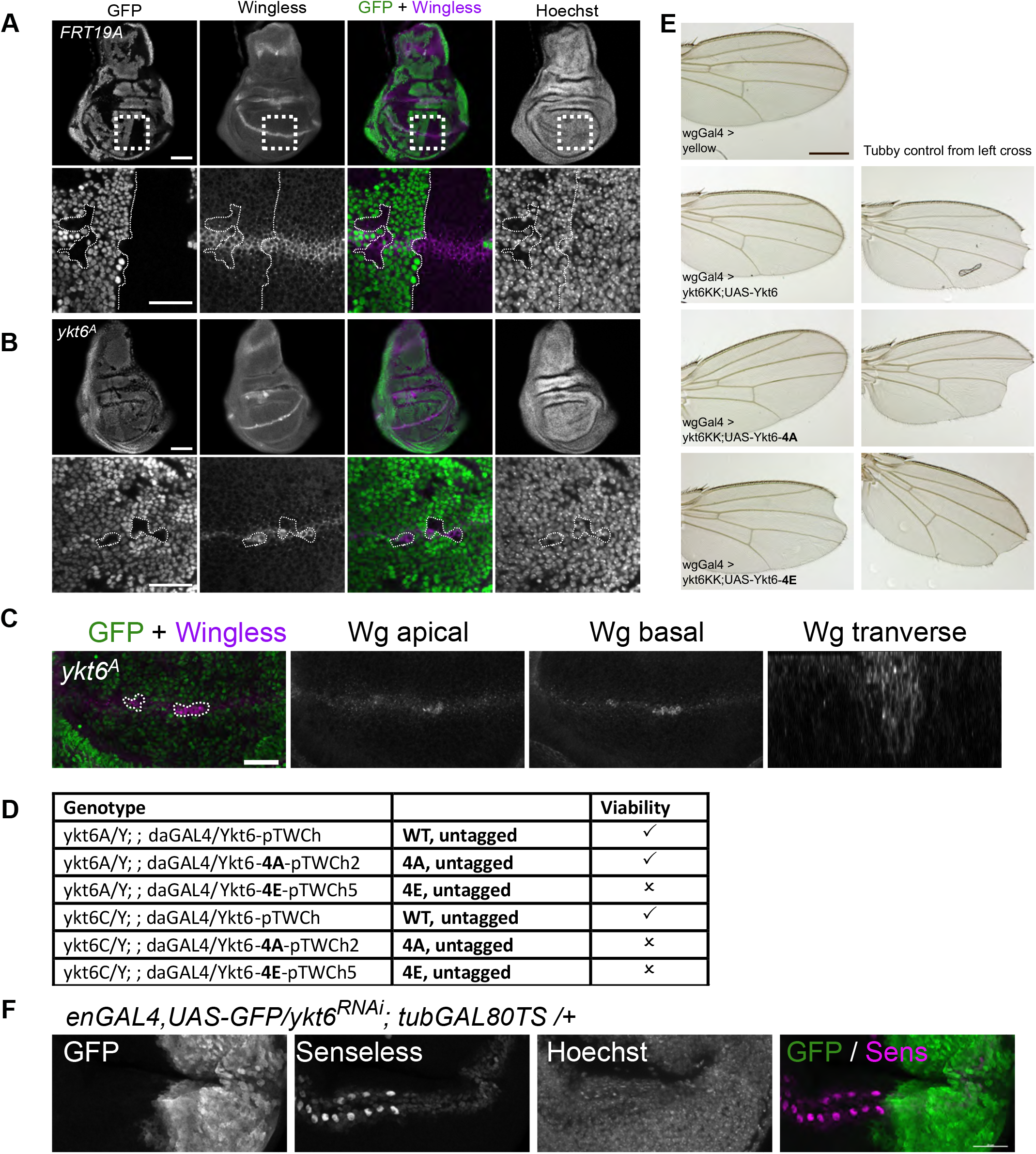
Ykt6 SNARE domain has an evolutionary conserved function in Wnt secretion. In contrast to FRT19A control clones (**A**) Wingless protein accumulates in *yktβ^A^* (**B,C**) clones marked by the absence of GFP. Panels in (A,B) show a merge of two subapical optical sections. Panels in (C) show a projection of five apical and five basal sections (distance 1 μm) as well as a transverse X-Z maximal intensity projection of the entire stack. Scale bars, 50 μm in overview and 20 μm in other images. (**D**) Rescue of lethality of ykt6^A^ and ykt6^C^ alleles. All Ykt6 constructs are inserted on the third chromosome (86Fb) and are expressed under the control of ubiquitous daughterless-GAL4. (**E**) Depletion of ykt6 by RNAi along the dorso-ventral border using wingless-GAL4 results in adult wing notches. Wing notches can be rescued by UAS-driven co-overexpression of WT and non-phosphorylatable Ykt6-4A but not phosphomimetic Ykt6-4E. All Ykt6 constructs are inserted on the third chromosome (86Fb). The images on the right show wings from the balancer flies from the same vial not co-expressing the UAS-Ykt6 constructs but only ykt6 RNAi as control. Scale bars 500 μm. These wings are representative of >10 wings from three independent experiments. (**F**) Time-controlled depletion of Ykt6 by RNAi (engrailed-Gal4, UAS-GFP/ UAS-ykt6RNAi; tubGal80-TS/+, larvae reared for three days at days at 29°C) causes reduction of Senseless staining in the posterior WID. Images are representative of > six WIDs from three independent experiments. Scale bar represents 20 μm.

**Fig. S2:**
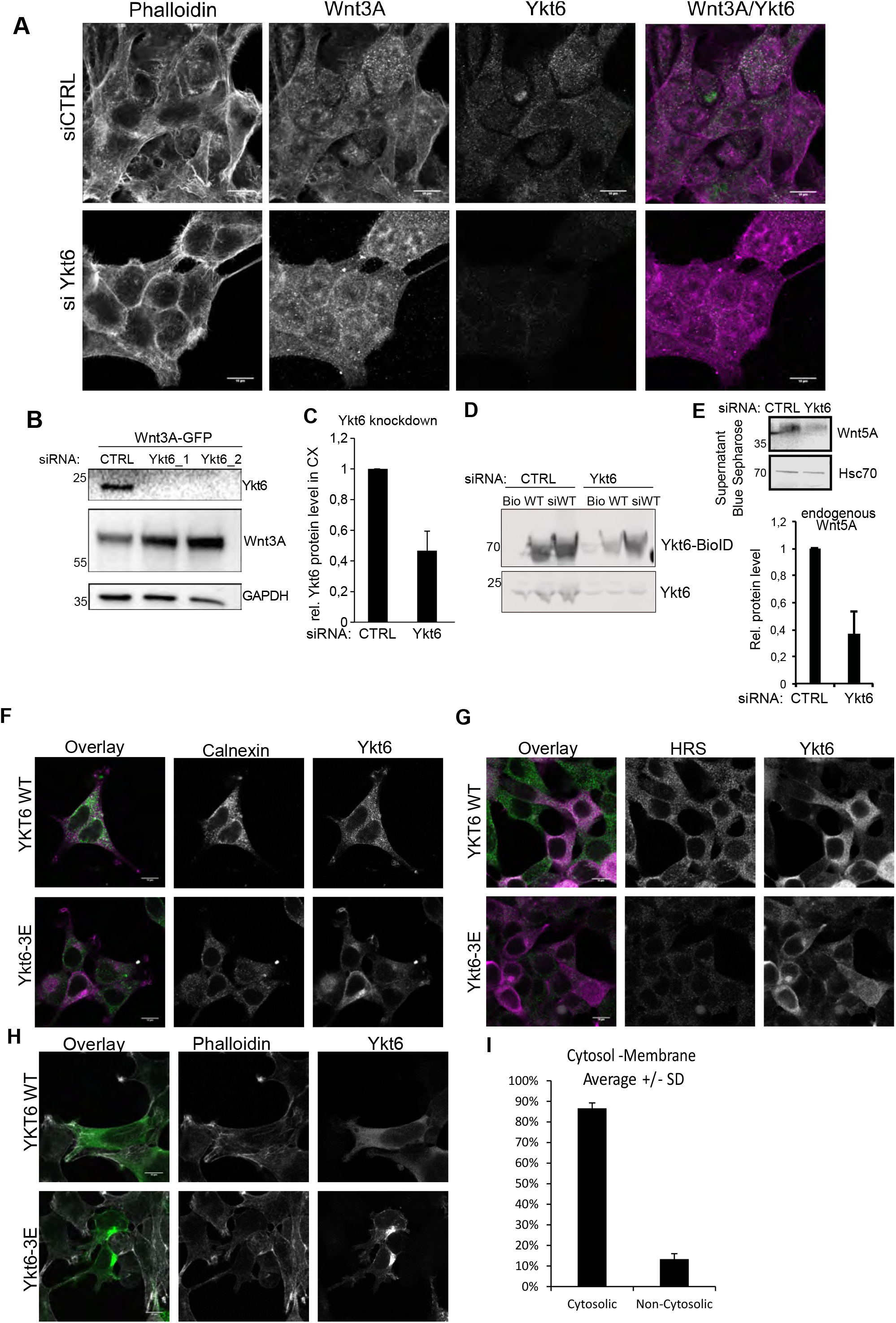
Wnt accumulates at post-Golgi membranes together with phosphomimicking Ykt6. (**A**) Confocal images of Wnt3A and Ykt6 in Hek293T cells with control and Ykt6 siRNA. (**B**) Wnt-GFP accumulates in Hek293T cells upon Ykt6 knockdown (**C**) Quantification of knockdown efficiency in Ykt6 siRNA1 and 2 transfected Hek293T cells from three independent experiments. (**D**) siRNA-resistant Ykt6-BioID construct in the presence of Ykt6 siRNA1 and 2. (**E**) Wnt5A secretion from SkBr3 cells is reduced in Ykt6 knockdown cells. Quantification of three independent experiments. (F-H) Colocalization of Bio-ID-tagged Ykt6-WT and -3E in Hek293T cells with organelle markers for ER, endosomes and plasma membrane: Calnexin (**F**), Hrs (**G**) and Phalloidin (**H**). (F-H) are representative images from three independent experiments, scale bars represent 10 μm. (**I**) Ykt6 quantitative proteomics data based on Organellar Maps from Itzhak *et al.*, 2016 places >80% of endogenous Ykt6 at the cytoplasm.

**Fig. S3.**
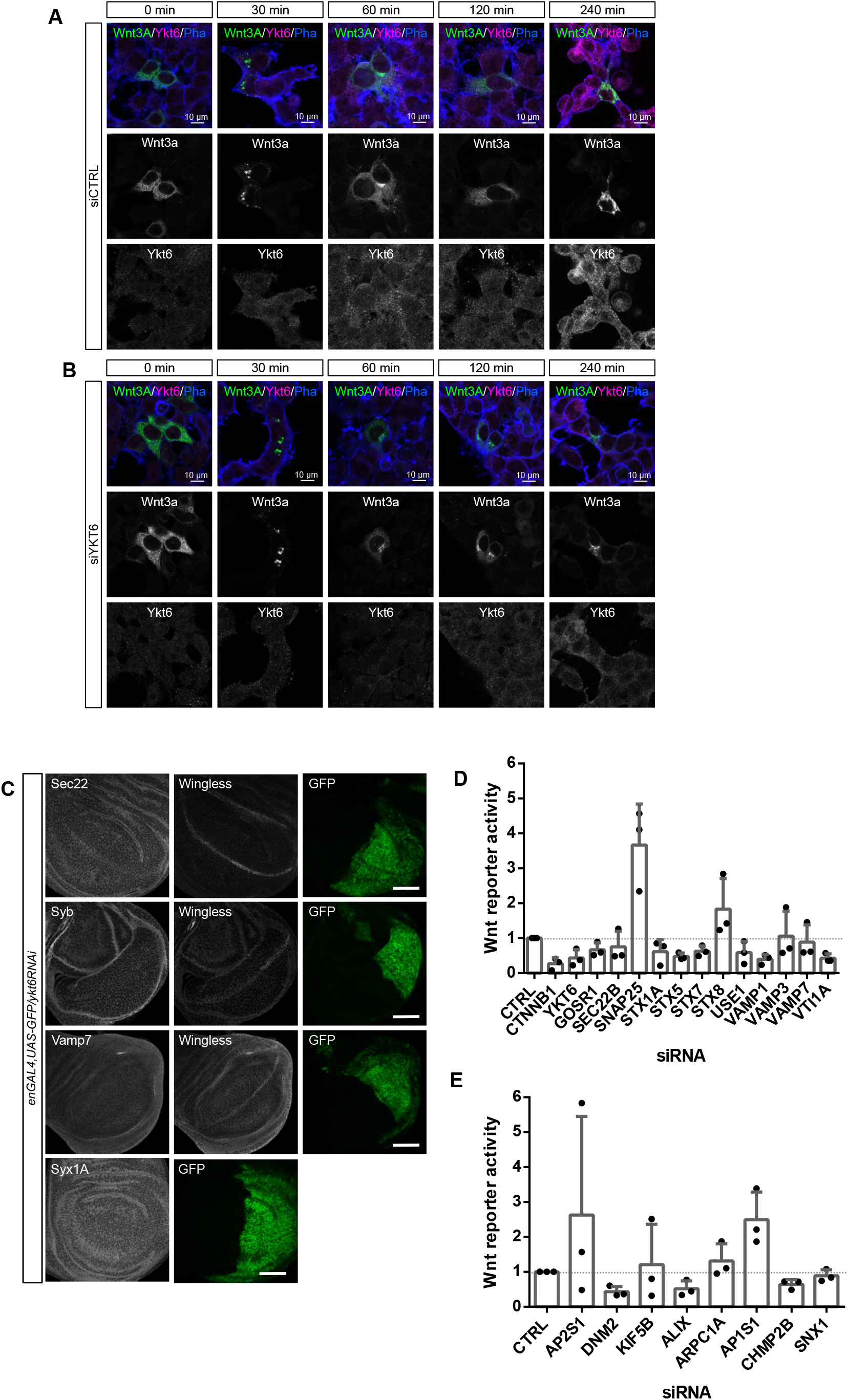
Mutant SNARE domain of Ykt6 blocks closed conformation. (**A**) Hek293T cells transfected with Rush-EGFP-Wnt3A together with control siRNA and (**B**) RNAi against Ykt6 were treated with 50 μM biotin for the indicated time points and stained for Wnt3A, Ykt6 and F-actin. (n=3), Scale bar represents 10 μm. (**C**) Knock-down of Ykt6 by RNAi in the posterior compartment of third instar WID marked by co-expression of GFP (engrailed-Gal4, UAS-GFP/UAS-ykt6RNAi) does not change the levels of Sec22, Syb, Vamp7 and Syx1A. Images in (C) are representative of > six WID per RNAi from two independent experiments. Scale bars represent 50 μm. (**D**) Wnt reporter assay of different SNAREs from three independent experiments. (**E**) Wnt reporter assay of endocytosis pathway components from Fig.3H from three independent experiments.

**Fig. S4:**
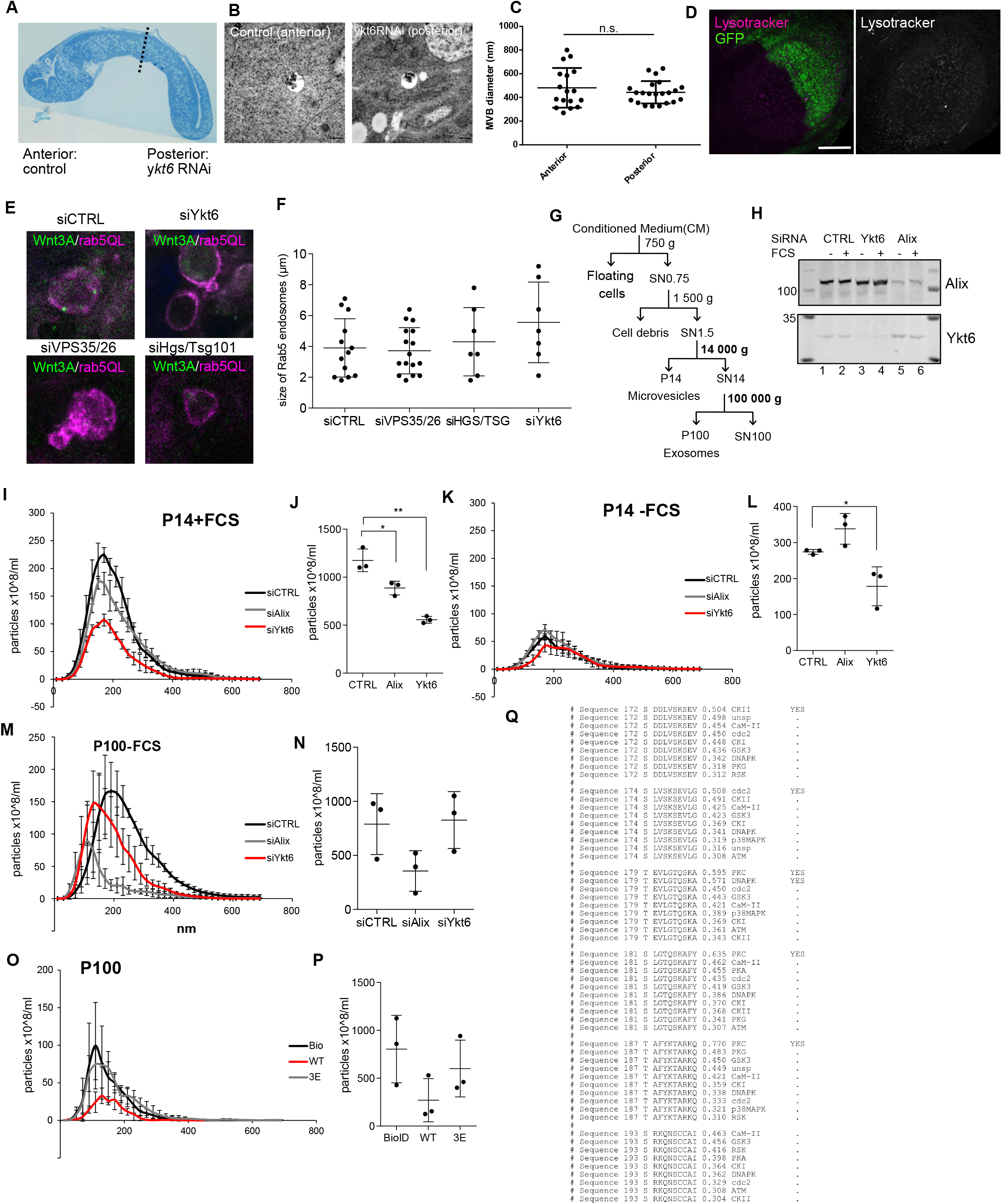
Ykt6 regulates exosome secretion in a concentration-dependent manner. (**A**) Semi-thin section of WID with time-controlled depletion of Ykt6 by RNAi (engrailed-Gal4, UAS-GFP/ UAS-ykt6RNAi; tubGal80-TS/+, larvae reared for three days at days at 29°C). (**B**) Electron microscopy images of WID. (**C**) Quantification of MVB size in electron microscopy images from cells in the anterior (n=17) and posterior (n=22) compartment of WID. (**D**) Knockdown of Ykt6 by RNAi in the posterior compartment of third instar WID marked by coexpression of GFP (engrailed-Gal4, UAS-GFP/UAS-ykt6RNAi) leads to increased Lysotracker staining. Images in (D) are representative of > six WID per RNAi from two independent experiments. Scale bar represents 50 μm. (**E**) Hek293T cells were co-transfected with plasmids for Rab5QL-DsRed and siRNA against Ykt6 (n=7), VPS26/35 (Retromer) (n=7) and HGS/Tsg101 (ESCRT) (n=16), or control siRNA (n=13) and analyzed by immunofluorescence microscopy. (**F**) Quantification of (E). (**G**) Differential centrifugation scheme of extracellular vesicles (EVs) purification. EVs in P14 are centrifuged 1h at 14,000g and EVs in P100 1h at 100,000g. (**H-N**) Secretion of EVs was analyzed by nanoparticle tracking analysis (NTA). (**H**) Western Blot of samples analyzed in (**I-N**) to control for knockdown efficiency. Size profile of P14-EV from normal (**I,J**) and starved (**K,L**) and of P100-EV from starved (**M, N**) Hek293T cells transfected with siRNA against control, Ykt6 or Alix. (**O,P**) NTA of P100 from Hek293T cells transfected with Ykt6-WT, -3E and BioID mock plasmid. (**Q**) PhosphoSite prediction for human Ykt6 using Netphos 3.1 (Blom *et al.*, 2004).

**Supplementary Table 1:** SNARE in vivo RNAi screening results. GAL4 virgin females were crossed to RNAi males as indicated. In a primary screen using wgGAL4, adult flies were scored for wing defects. RNAi lines producing adult wing notches were further verified for Wg secretion defects in larval WID using enGAL4,UAS-GFP.

| Gene name | Category |  | Human homologue | Lines | wgGAL4 X RNAi | enGAL4,UAS-GFP X RNAi | References |
| --- | --- | --- | --- | --- | --- | --- | --- |
| **Bet1** | Qc | t | Bet1 | GD8421 | **Notches** | **Lethal** |  |
| Gos28 | Qb | t | GOSR1 (Golgi SNAP receptor1) | KK100289 | No phenotype |  |  |
| **Membrin** | Qb | t | GOSR2 (Golgi SNAP receptor2) | KK109404 | **Notches** | **Mild effect** |  |
| nSyb | R | v | Vamp2, neuronal SNARE | BL31983 | n.a. |  |  |
| **Sec20** | Qb | t | BNIP1, SEC20 | KK100264 | **Notches** | **No effect** |  |
| **Sec22** | R | v | Sec22b | BL34893 | **Notches** | **Mild effect** | Li, BBRC 2015 |
| **SNAP24** | Qbc | t | SNAP25 | KK108209 | **Notches** | **No effect** |  |
| SNAP25 | Qbc | t | no homologue | TRiP27306, 34377 | No phenotype |  |  |
| SNAP29 | Qbc | t | SNAP29 | KK107947 | No phenotype |  |  |
| **Syb** | R | v | Vamp1 | KK102922 | **Notches** | **Wg intracellular accumulation** | Yamazaki, NCB 2016 |
| **Syx1a** | Qa | t | STX1A | GD33112 | **Notches** | **Lethal** |  |
| Syx4 | Qa | t | no homologue | n.a. | n.a. |  |  |
| **Syx5** | Qa | t | STX5 | KK108928 | **Notches** | **Lethal** |  |
| **Syx6** | Qc | t | STX6 | KK104795 | **Notches** | **No effect** |  |
| **syx7/avl** | Qa | t | STX7 | KK107264 | **Lethal** | **Lethal** |  |
| **Syx8** | Qc | t | STX8 | KK107014 | **Notches** | **Mild effect** |  |
| **Syx13** | Qa | t | TSNARE1 | KK102432 | **Notches** | **No effect** |  |
| Syx16 | Qa | t | STX16 | KK109504 | No phenotype |  |  |
| Syx17 | Qa | t | STX17 | KK108825 | No phenotype |  |  |
| **Syx18** | Qa | t | STX18 | KK105113 | **Notches** | **Lethal** |  |
| **Use1** | Qc | v | USE1 | KK100019 | **Notches** | **Wg intracellular accumulation** |  |
| Vamp7 | R | v | VAMP7 | KK108733 | No phenotype |  | Gao, Plos One 2017 |
| **Vti1a** | Qb | t | VTI1A | KK109819, | **Notches** | **Wg reduction** |  |
| Vti1b | Qb | t | no homologue | n.a. | n.a. |  |  |
| **Ykt6** | R | v | YKT6 | KK1515 | **Notches** | **Lethal** | Gross, NCB, 2012 |

**Supplementary Table 2:**
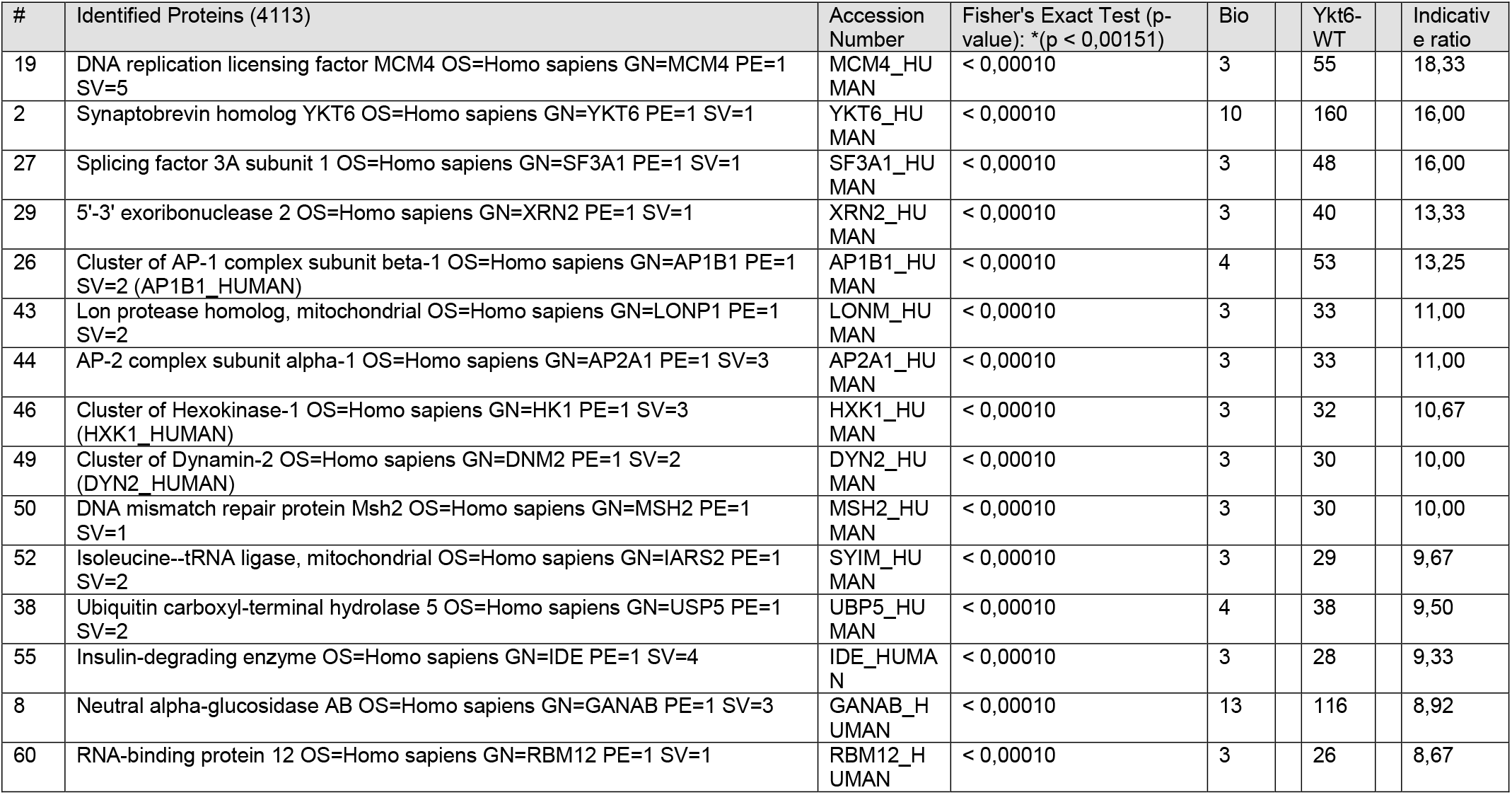
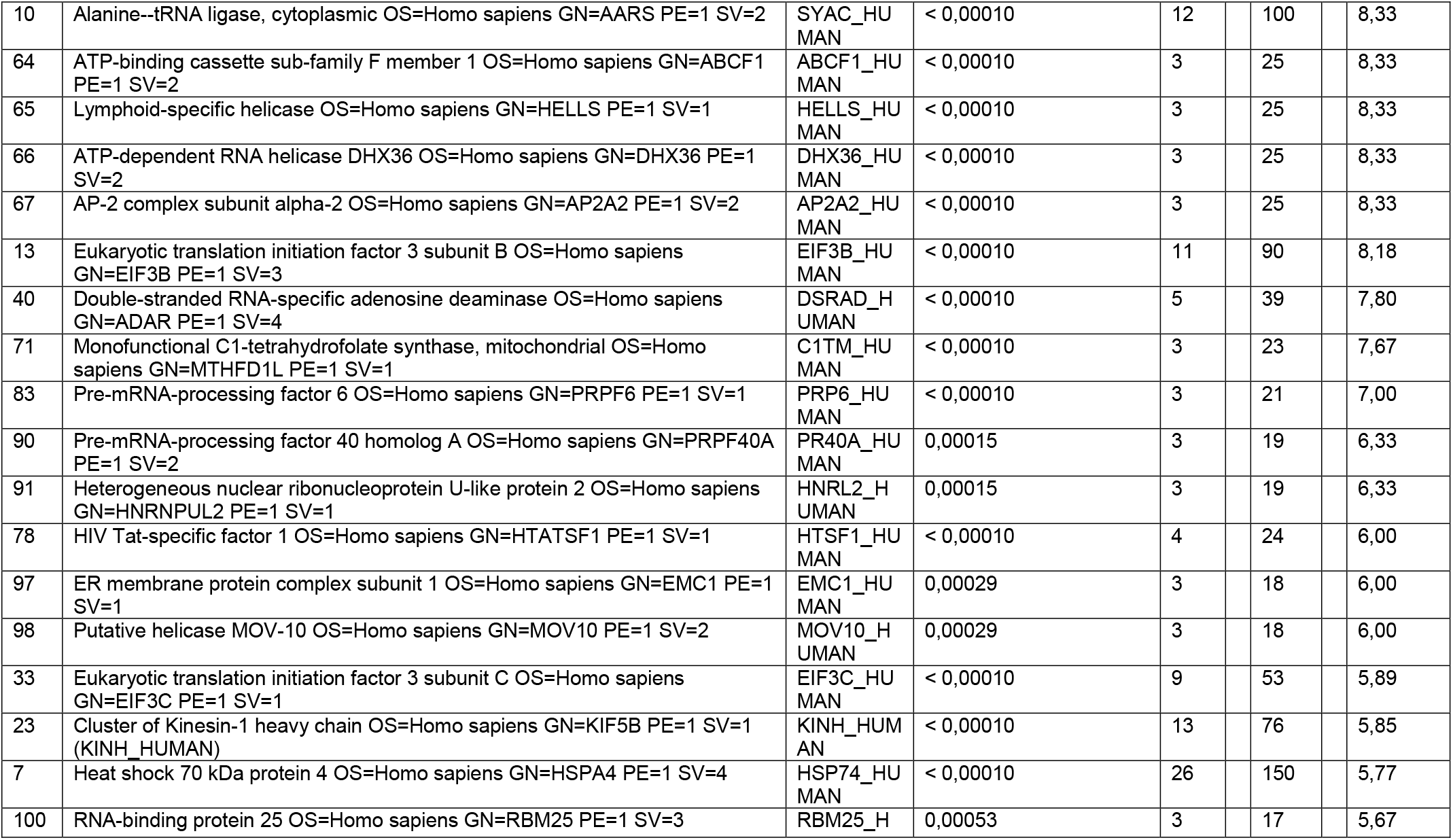
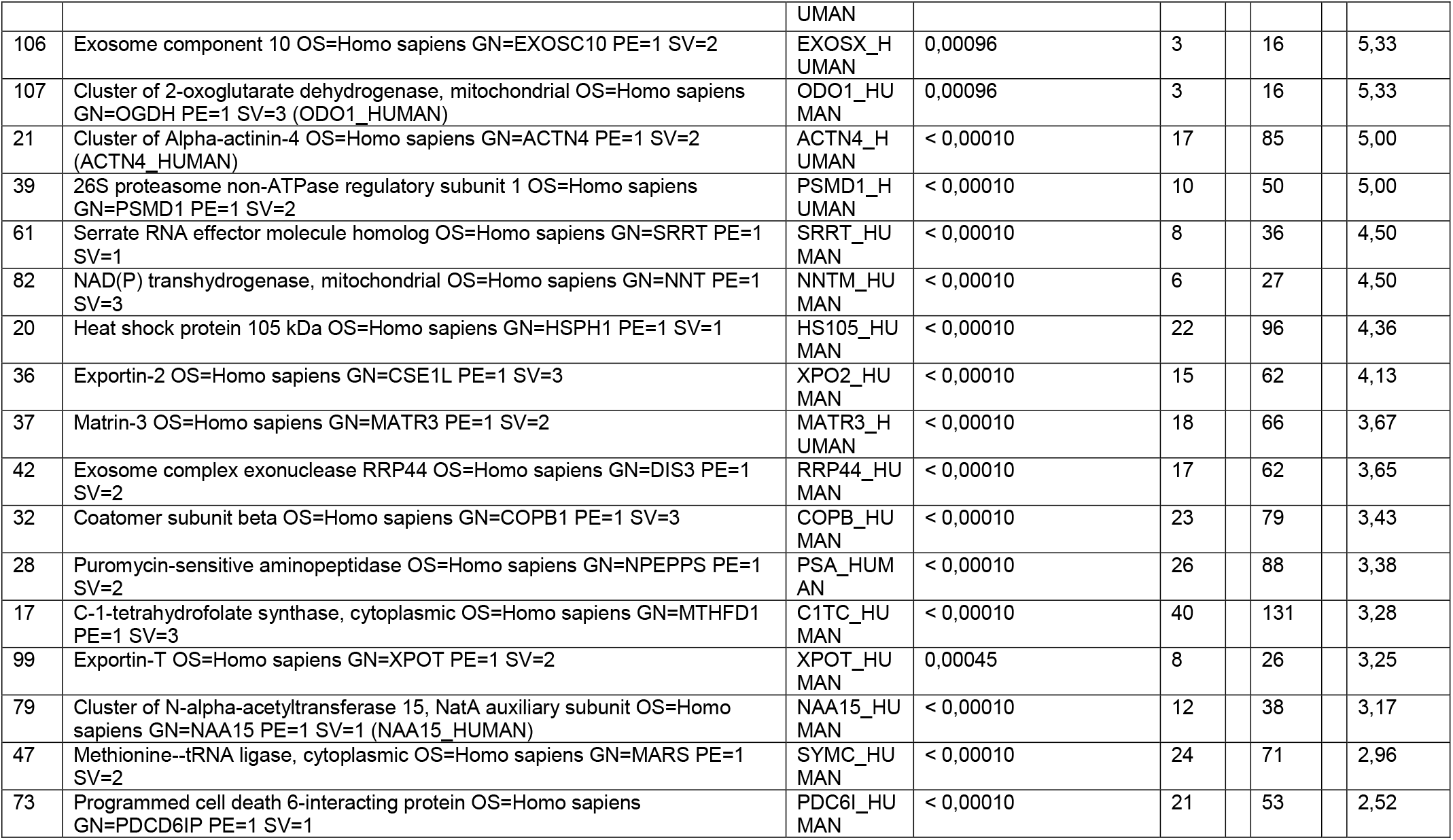
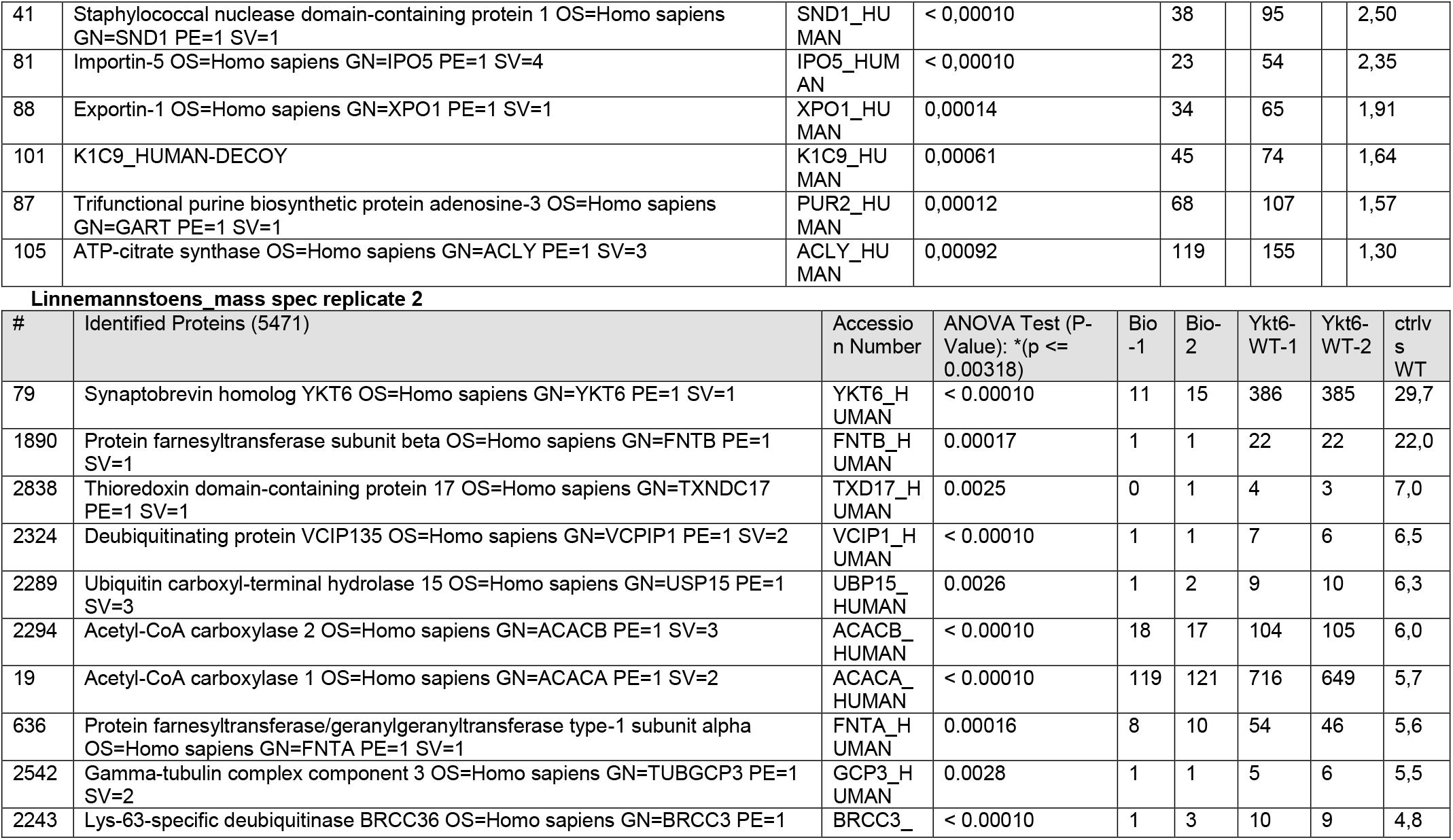
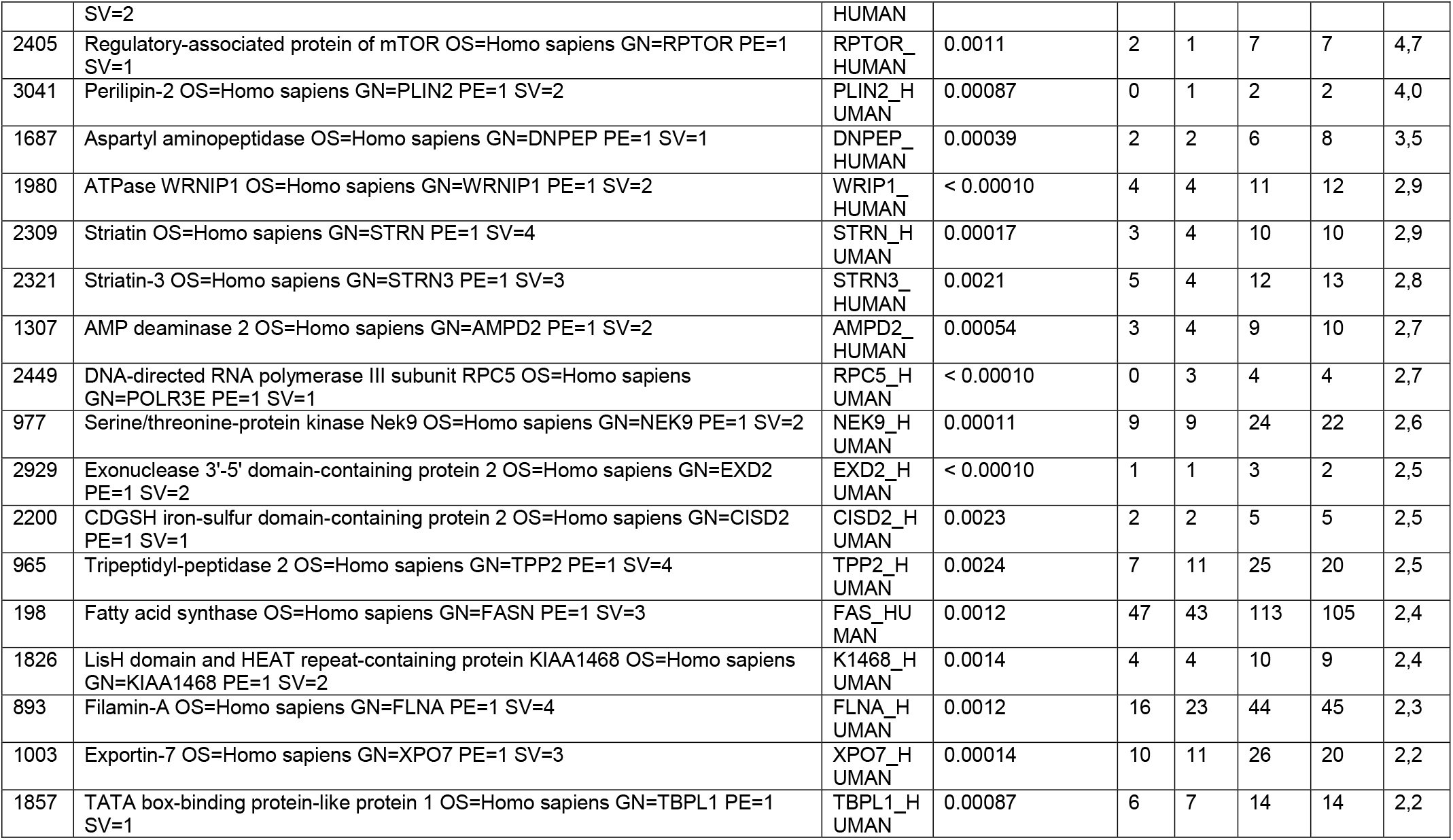
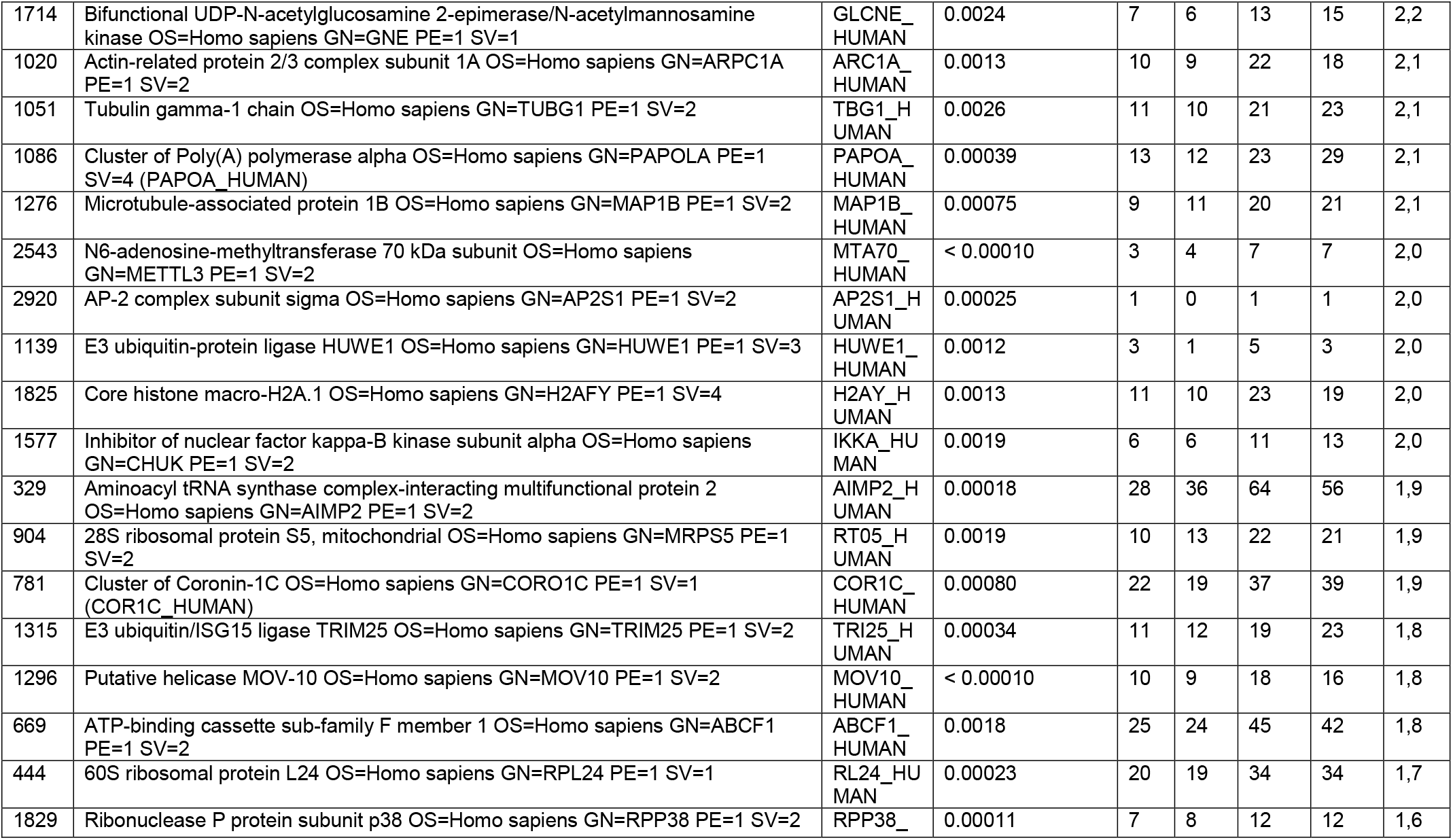
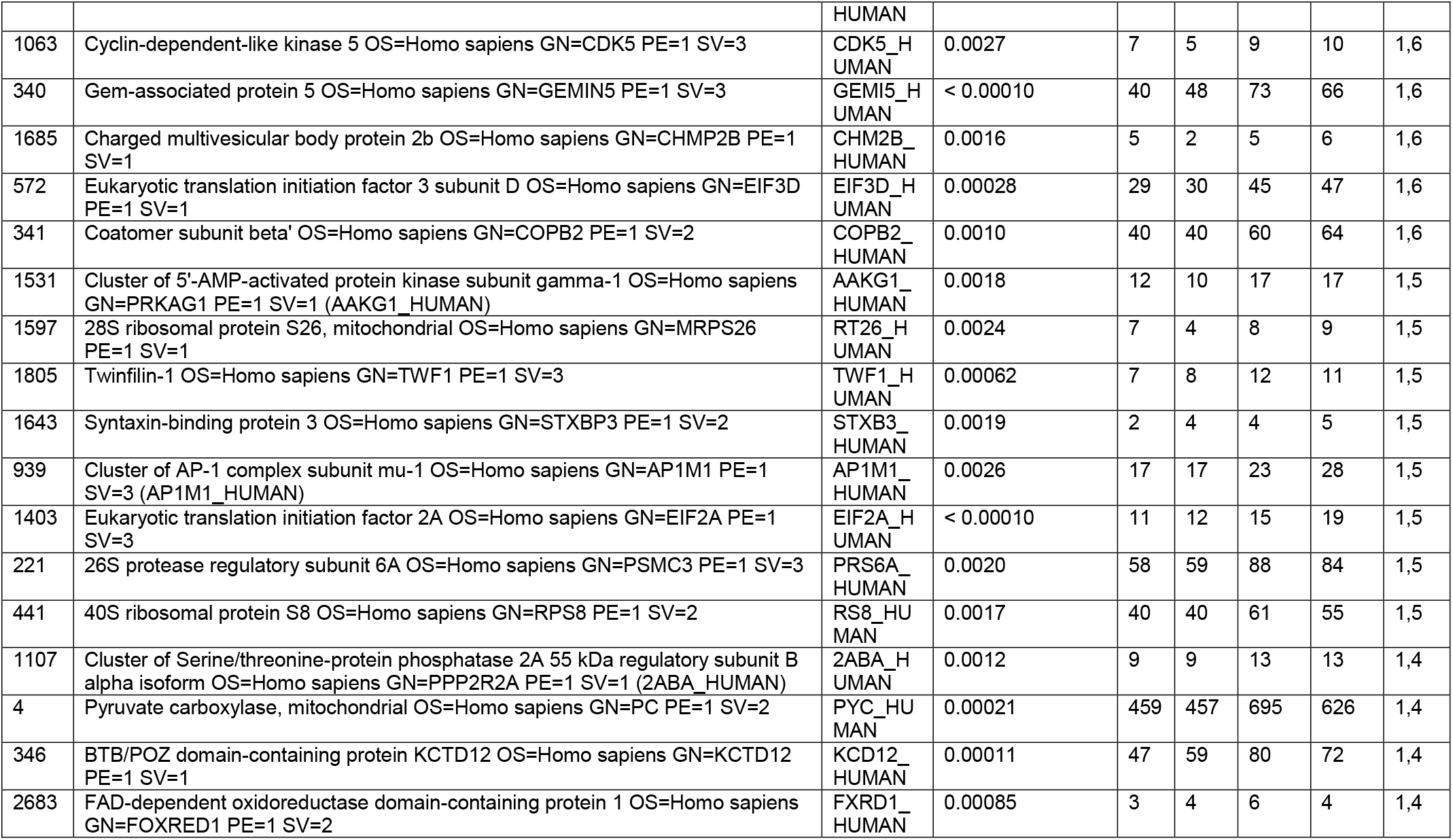
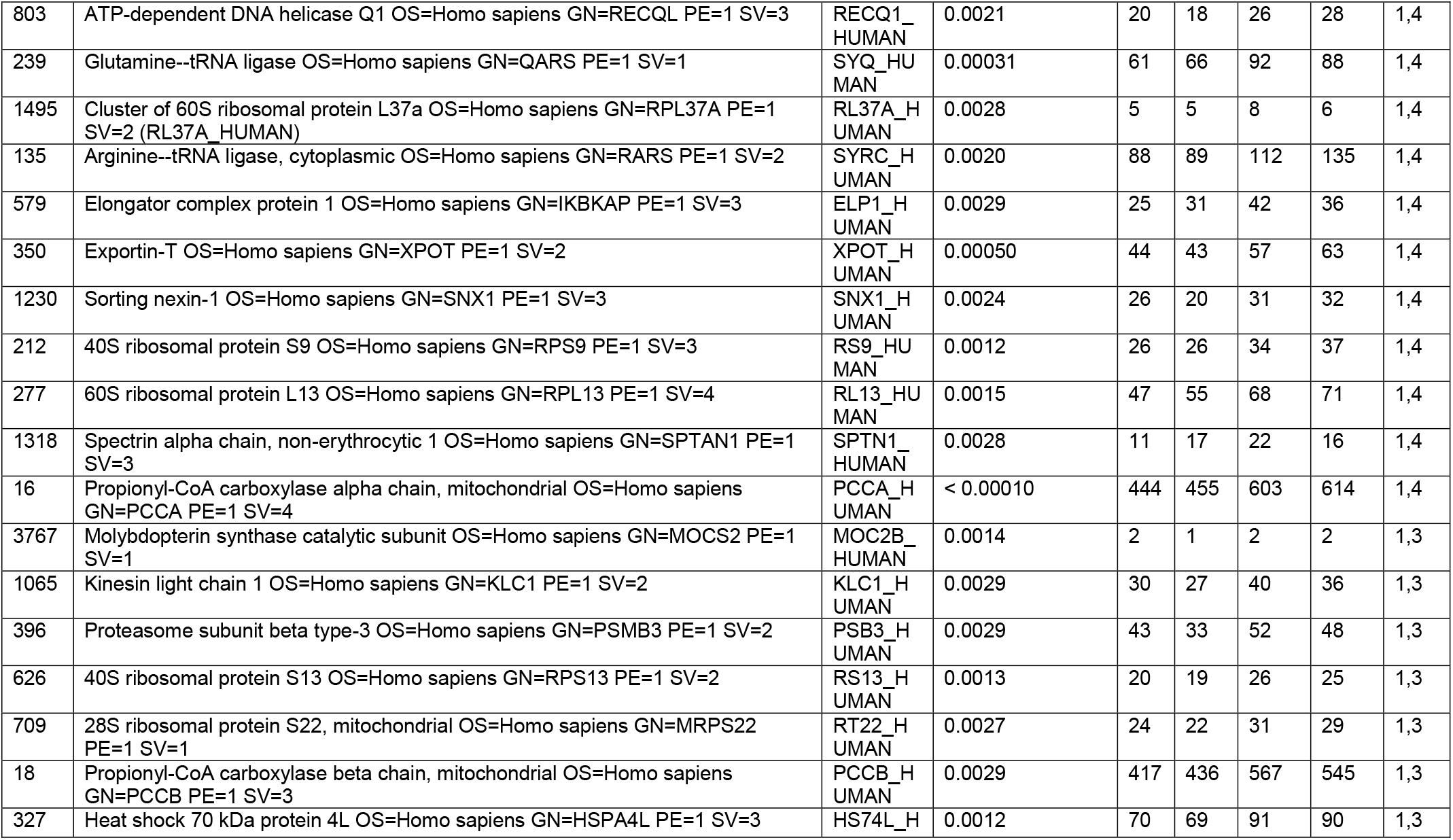
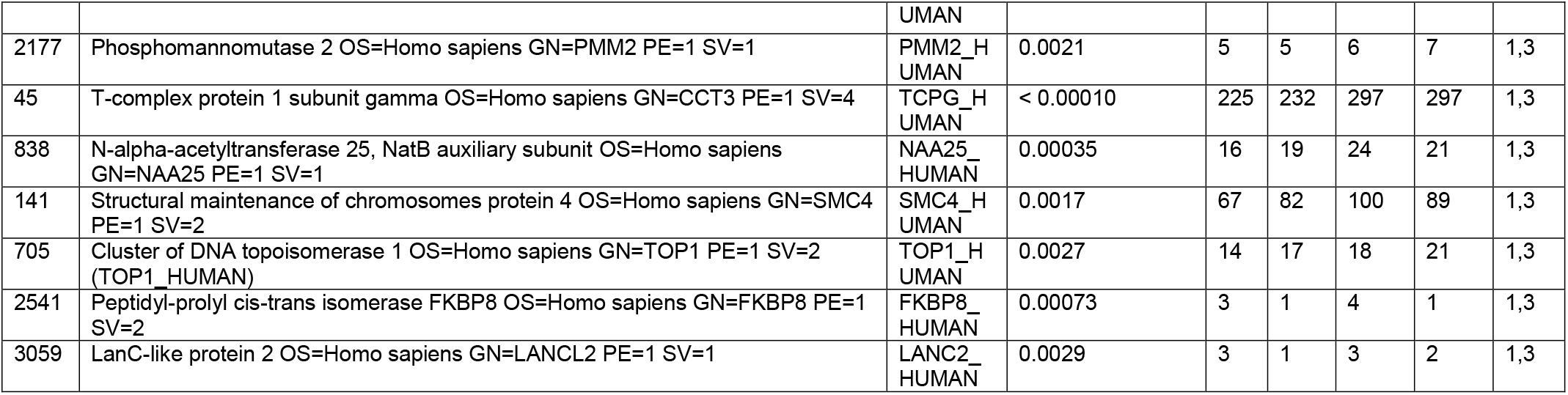
Proteins identified by mass spectrometry from BioID control and Ykt6-WT samples in two biological replicates, replicate_1 by SWATH-MS analysis, replicate_2 by Spectral Counting analysis (for experimental details see Material and Methods section)

**Supplementary Video 1:** Time-lapse movie of RUSH-EGFP-Wnt3A in Hek293T cells was recorded for 4h 28min with 60 sec intervals at 37°C after addition of 50 μM biotin. Scale bar 20μm. Movie exported at 10fps.

